# Effect of Immunosuppressive Drugs on Glucose-Stimulated Insulin Secretion: Concentration-Response Studies in Dynamic Perifusion Assays

**DOI:** 10.64898/2026.07.09.737557

**Authors:** Sung-Ting Chuang, Brandon Watts, Oscar Alcazar, Peter Buchwald

## Abstract

Immunosuppressive drugs, which are required to maintain graft function in transplant recipients, are associated with many unavoidable side effects including posttransplant diabetes mellitus (PTDM) that involves both peripheral insulin resistance and impairment of insulin secretion. To characterize in detail the concentration-dependency of the effect of well-known immunosuppressive drugs on glucose-stimulated insulin secretion (GSIS), we performed dynamic perifusion studies with human pancreatic islets. The effect on the time-profile of GSIS has been assessed over a wide concentration range for several clinically relevant immunomodulatory therapies, including small-molecule drugs (cyclosporine, sirolimus, tacrolimus, prednisolone acetate, and loteprednol etabonate) and biologics (abatacept and anti-CD40L), plus a prospective β-cell proliferation-inducing agent (harmine). While biologics showed no significant detrimental effects after one-day treatment even at relatively high concentrations (5 µM), all small-molecule drugs inhibited insulin secretion in a concentration-dependent manner, although glucocorticoids showed a distinct response pattern. Calcineurin and mTOR inhibitors preserved GSIS within their therapeutic ranges but progressively distorted its time-profile at higher concentrations and completely suppressed secretion at the highest levels. Cyclosporine exhibited the least, only about 35-fold, separation between its therapeutic target (*C*_targ_) and half-maximal GSIS inhibitory (IC_50_) concentrations. Glucocorticoids did not alter the shape of the time-profile but inhibited overall insulin secretion even at therapeutic levels. Their inhibitory effect only increased slowly with concentration and did not follow a classic sigmoid pattern that has unity Hill slope. These findings establish quantitative benchmarks for immunosuppressant-induced β-cell toxicity and provide a framework for optimizing immunosuppressive regimens to reduce the risk of PTDM.

## Introduction

The therapeutic promise and widespread applicability of organ and cell transplantation is considerably limited by the need for lifelong immunosuppression to prevent rejection because systemic and long-term immunosuppression with all currently available agents is unavoidably accompanied by serious side effects. These include renal toxicity, cardiovascular morbidity, and increased susceptibility to infections and malignancies (cancer), among others, especially if higher treatment doses are required [1]. Side effects also include posttransplant diabetes mellitus (PTDM), which can occur in up to 40–50% of patients and is largely driven by the immunosuppressive therapy, particularly corticosteroids, calcineurin inhibitors, and mTOR inhibitors [2–4]. Recipients of solid organ transplant have a nine-fold increased risk of diabetes compared to age-matched controls [3]. The pathogenesis of PTDM involves both peripheral insulin resistance and dysfunction of β-cells (impaired insulin secretion) [2, 4].

These problems are further compounded in the case of β-cell replacement / islet cell transplantation for treatment of type 1 diabetes (T1D) [5–7] as all existing systemic immunosuppression regimens have serious negative effects on the engraftment, function, and survival of transplanted islets, thereby limiting the success of these therapies [8, 9]. Due to our interest in immunotherapies for β-cell replacement / islet cell transplantation therapies including via localized delivery [10, 11], we are particularly interested in characterizing the effect of immunosuppressive drugs on the insulin secreting ability of human islets. Accordingly, here we studied several clinically relevant immunomodulatory therapies and investigated the concentration dependence of their effects on glucose-stimulated insulin secretion (GSIS) of human islets using dynamic perifusion studies. Among small-molecule drugs, which are known to hamper insulin secreting ability especially at higher concentrations, we evaluated the effects of two calcineurin inhibitors (CNIs; cyclosporine and tacrolimus), the mTOR inhibitor sirolimus (rapamycin), two glucocorticoids (prednisolone acetate and loteprednol etabonate), and a prospective β-cell proliferation-inducing agent (harmine). Among biologics, which are less likely to affect GSIS, we evaluated CTLA4-Ig (abatacept) and anti-CD40L (5C8H1).

Interestingly, while it is widely recognized that most immunosuppressive drugs have deleterious effects on insulin secretion, which as any pharmacological response has to be concentration-dependent, there are very few, if any, detailed studies characterizing this in general and especially using dynamic perifusion studies and human islets. Perifusion studies, which assess the detailed time-profile of insulin secretion in response to a high glucose or other secretagogue challenge, are the most complex *in vitro* assay that can be used to assess the quality and function of isolated islets or other insulin secreting cells [12]. They have been around for more than 50 years [13–15] and due to considerable technical and analytical advances since then, they allow the detailed dynamic characterization of GSIS under controllable influxes of glucose, oxygen, and other agents of interest. They provide much more information-rich characterization than the more widely used static GSIS as they use flowing media allowing better oxygenation and thus function of the insulin secreting cells and can assess both first- and second-phase insulin secretion [16, 17]. Because islets are mini organs, they can function quite well even after isolation and produce responses similar to those in their original physiological environment. Therefore, *ex vivo* perifusion studies with isolated islets can be used to characterize in detail responses that otherwise would be difficult to obtain *in vivo*.

## Methods

### Materials

Immunomodulatory drugs and biologics were obtained from commercial sources and used as such; they were as follows: cyclosporine (cat. no. 300152), sirolimus (rapamycin; 100766), tacrolimus (100795), and harmine (329605) from MedKoo (Durham, NC, USA); loteprednol etabonate (SML0547) from Sigma-Aldrich (St. Louis, MO, USA); prednisolone acetate ophthalmic suspension (NDC 61314-637-10) from Sandoz (Princeton, NJ, USA); loteprednol etabonate ophthalmic suspension as Lotemax® (NDC 82260-299-05) from Bausch & Lomb (Bridgewater, NJ, USA); abatacept from Bristol Myers Squibb (New York, NY, USA); anti-CD40L (5C8H1, PR-1547, RRID: AB_2716324) from NHPRR (Nonhuman Primate Reagent Resource; Boston, MA, USA). Glucose, NaCl, KCl, CaCl_2_, MgCl_2_, HEPES, and bovine serum albumin (BSA) from Sigma-Aldrich (St. Louis, MO, USA); islet culture media from Prodo Laboratories (Aliso Viejo, CA, USA).

### Human islets

Human pancreatic islet samples were procured from the Integrated Islet Distribution Program (IIDP; RRID:SCR_014387) at City of Hope (Duarte, CA, USA). All islet samples used here were from non-diabetic donors; characteristics of the human islet donors for the present study are summarized using standard checklists recommended for reporting human islet preparations used in research in Supplementary Material Table S1.

### Islet perifusion

Islet samples aliquoted into batches of around 300 islet equivalents (IEQ) were maintained under standard culture conditions of 37°C in standard islet culture media (Prodo Laboratories, Aliso Viejo, CA, USA) in the presence of various concentrations of test drugs for 24 h. The perifusion experiments (dynamic GSIS) were performed using a PERI4 machine (Biorep Technologies, Miami, FL, USA; RRID:SCR_004907) that allows parallel perifusion of up to 12 channels via a microfluidic manifold. For each experiment, an estimated 100 IEQ of human islets were handpicked and loaded into Perspex microcolumns between two layers of acrylamide-based microbead slurry (Bio-Gel P-4, Bio-Rad Laboratories, Hercules, CA, USA) by experienced operators. Perifusion buffer containing 125 mM NaCl, 5.9 mM KCl, 1.28 mM CaCl_2_, 1.2 mM MgCl_2_, 25 mM HEPES, and 0.1% bovine serum albumin (BSA), adjusted to pH 7.4 and maintained at 37°C with selected glucose or KCl (25 mM) concentrations was flowed through the columns at a rate of 100 μL/min. After 1 h of washing with low glucose solution (4 mM, G4) for stabilization, islets were exposed to the following sequence for which the outflow was collected: 8 min of low glucose (G4), 20 min of high glucose (16 mM, G16), 15 min of low glucose (G4), 10 min of KCl, and 10 min of low glucose (G4). Samples (100 μL) were collected every minute from the outflow tubing of the columns in an automatic fraction collector designed for a multi-well plate format. The islets and the perifusion solutions were kept at 37°C in a built-in temperature-controlled chamber while the perifusate in the collecting plate was kept at <4°C to preserve the integrity of the analytes. Insulin concentrations were determined with commercially available human ELISA kits (Mercodia Inc., Winston-Salem, NC, USA). Values obtained with the human kit were converted from mU/L to μg/L using 1 μg/L = 23 mU/L per the manufacturer guidelines. Here we used a G4–G16–G4 low–high–low glucose sequence instead of the previously used G3– G11–G3 one because our previous data suggested it as better suited to characterize the GSIS of human islets [16]. Normalization of the insulin secretion to DNA, protein, or insulin content was not used [18]. However, to account for possible differences among the islet mass loaded into the different perifusion channels, as accurately assessing it in IEQ units is nontrivial [19, 20], values were adjusted based on the response to KCl using the area under the curve (AUC) in the control (untreated) columns for normalization as described before [16, 21–23]. All responses were scaled to 100 IEQ. Average perifusion responses to such stepwise low–high–low glucose challenges shown for comparison are from our previous work done following the same procedure as described there [16, 17]. Islet insulin content was determined by incubating the islet pellet retrieved from the perifusion chamber in 200 μL of fresh acid ethanol (50 μL 5N HCl + 5.5 mL 95% ethanol) for 24 h [24], which was then analyzed using the same ELISA kit used for insulin concentration determination.

### Statistical analysis

Data used here are averages of at least three samples for each condition. All graphic preparations and curve fittings were performed using GraphPad Prism 11 (GraphPad, La Jolla, CA, USA; RRID:SCR_002798). Half-maximal inhibitory concentration (IC_50_) values for the AUC data were obtained using nonlinear regression with the log(inhibitor) vs normalized response model in Prism.

## Results

To study in detail the effect of immunosuppressive drugs on glucose-stimulated insulin secretion, we performed dynamic perifusion studies with human pancreatic islets pretreated with increasing concentrations of therapeutics currently approved for immunomodulatory therapies. By using a perifusion-dedicated automated machine (PERI4) with up to 12 channels and outflow collection with adjustable temporal resolution (1 min used here), islets from the same batch exposed to different conditions could be perifused in parallel to allow a direct comparison of the differences in the responses. This way, we could assess in detail the concentration-dependent effect on the time profile of insulin response from the same islets, i.e., effects on the first- and second-phase insulin releases as well as on the total insulin released. We studied several clinically relevant immunomodulatory small-molecule drugs (cyclosporine, sirolimus, tacrolimus, prednisolone acetate, and loteprednol etabonate) and biologics [CTLA4-Ig (abatacept) and anti-CD40L (5C8H1)], plus a prospective β-cell proliferation-inducing agent (harmine) of interest in regenerating the insulin-producing β-cells that are destroyed in T1D.

All studies presented here were performed with a classic low–high–low glucose step (“square wave”). Here, we used a 4 – 16 – 4 mM (G4–G16–G4) sequence – slightly different from our previously used G3–G11–G3 step because our earlier studies of the concentration-response of glucose-stimulated insulin secretion suggested this as better suited to characterize the GSIS of human islets [16]. Use of this step had no significant effect on the general shape of the time-profile of insulin release induced by a square wave of low-high-low glucose step as illustrated in Figure 1 that shows the average profiles obtained for control untreated islets obtained here with this G4–G16–G4 step (*n* = 47) compared to those obtained by us earlier with the G3–G11–G3 step (*n* = 55). For additional reference, the time-profiles previously published by us (*n* = 34 [16] and *n* = 70 [17]) are also included in Figure 1 together with another one obtained by a different group with a different equipment and perifusion protocol as part of the assessment for the Integrated Islet Distribution Program (IIDP), the main source of human islets for research within the US (G5.6–G16.7–G5.6, *n* = 299) [25]. The average profile obtained by us remained essentially unchanged despite the change to the current G4–G16–G4 glucose step: it was somewhat lowered in general due to the improved methodology and perifusion apparatus reducing the stress on the islets, a classic cause of elevated baseline secretion. Even the average profile obtained by IIDP with a different protocol and different equipment is quite similar just showing larger variability and somewhat more elevated secretion (data from [25] rescaled to one IEQ and with two of the last high-glucose points omitted as a longer exposure time of 30 min was used in that study) (Figure 1).

**Figure 1.**
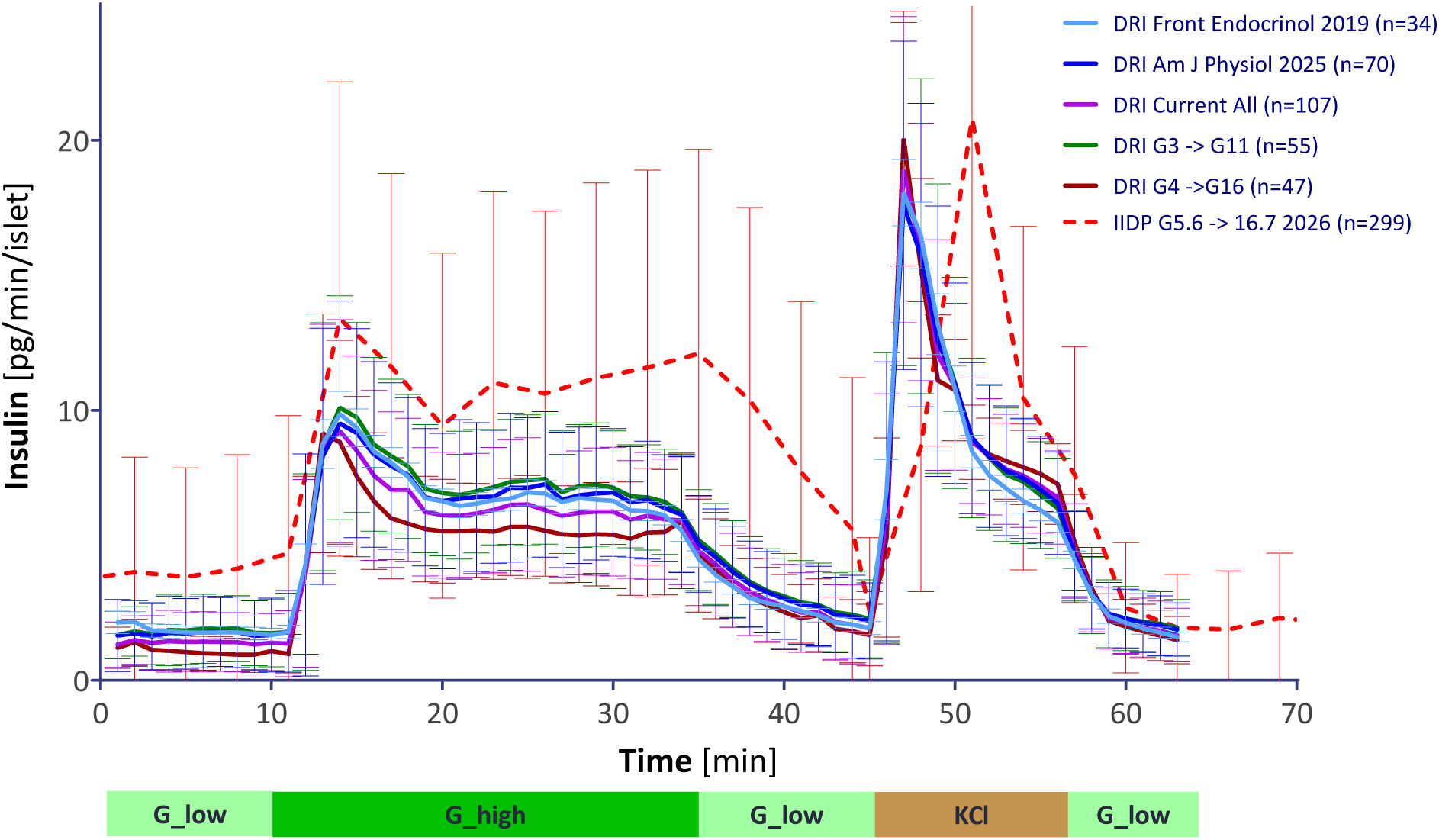
Average time profiles of insulin secretion rates of perifused human islets (dynamic GSIS). Average time-profile of untreated control islets perifused using our automated multichannel perifusion apparatus (PERI4) and the current protocol (G4→G16: low G4 for 8 min, high G16 for 20 min, low G4 for 15 min glucose stimulation followed by 25 mM KCl and G4, each for 10 min; *n* = 47; dark red) versus that obtained with our previous protocol (G3→G11; *n* = 55; green). For comparison, two time profiles previously published by us (*n* = 34 [16] and *n* = 70 [17]; light and dark blue, respectively) are also included together with another one obtained by a different group (IIDP) with a different protocol on a large number of islets (G5.6→G16.7; *n* = 299; red) [25]. Data shown are average ± SD with the number of samples (*n*) as indicated in the graph.

### Calcineurin inhibitors – cyclosporine

Cyclosporine (cyclosporin A; Sandimmune®), a cyclic undecapeptide CNI, was approved by the FDA in 1983 and significantly expanded the applicability of organ transplantation. Cyclosporine (CsA) is used to treat organ rejection post-transplant as well as in certain autoimmune diseases, such as rheumatoid arthritis (RA), psoriasis, and amyotrophic lateral sclerosis (ALS). It is also used as an eye drop in keratoconjunctivitis sicca (dry eye). Its therapeutic range in whole blood is 100–500 ng/mL (83–415 nM) depending on the application, and CsA is known to have a relatively narrow therapeutic index (separation between effective and toxic concentrations) [26–28]. Here, we explored its effect on dynamic GSIS at concentrations ranging from 40 to 25,000 nM using five-fold increments (Figure 2). As clearly noticeable in the figure, concentrations slightly higher than the therapeutic target (*C*_targ_ ≈ 300 nM) and probably within the range of peak concentrations (*C*_max_) are already affecting insulin secretion. First-phase secretion (the peak during the first 5–10 min following high-glucose exposure) disappears and total insulin secretion is already diminished at 5,000 nM (5 μM; Figure 2, top). The highest concentration tested (25 μM) considerably diminished GSIS, reducing it by 80% without significantly affecting KCl-induced insulin release. Fitting of the area under the curve (AUC_ins_) data, a measure of total insulin secreted, with a classic sigmoid concentration-response function suggested an estimated half-maximal inhibitory concentration of IC_50_ = 10,800 nM (Figure 2, bottom) – only an about 35-fold separation compared to *C*_targ_ (Table 1). Here, we use target concentration, *C*_targ_, as the estimated concentration that has to be reached and maintained to achieve therapeutic effects while minimizing the adverse effects (AEs); this, of course, depends somewhat on the specific application. We selected *C*_targ_ as 300 nM for CsA based on literature data [26, 27].

**Figure 2.**
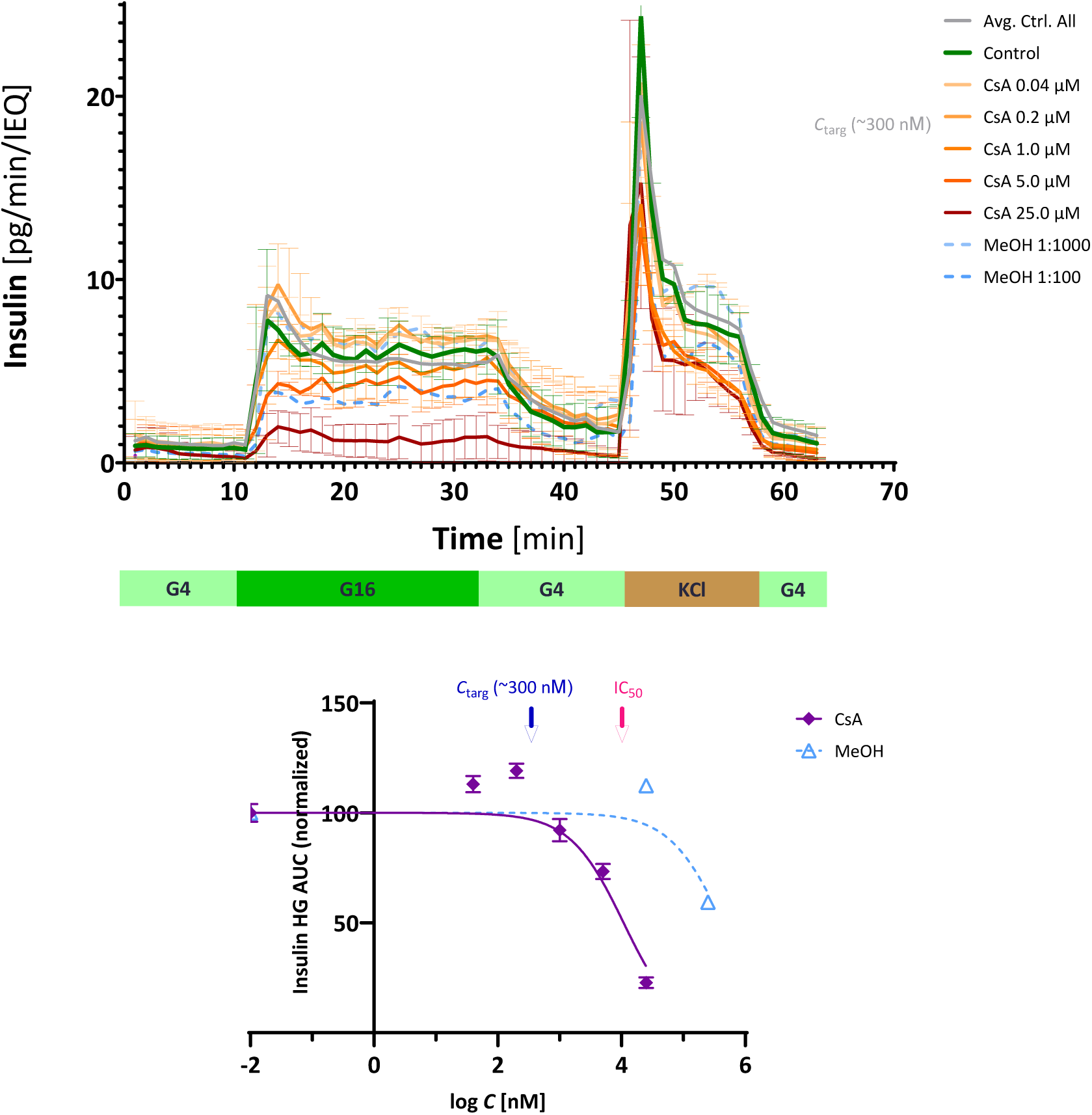
Concentration-dependence of the effect of cyclosporine on the GSIS of human islets. (Top) Time-profile of glucose-stimulated insulin secretion (GSIS) of isolated islets exposed to increasing concentrations of cyclosporine (CsA) for 24 h and then perifused using a stepwise sequence of low-high-low glucose (G4–G16–G4) as indicated. Increasing CsA concentrations are denoted with increasingly darker colors; the methanol (MeOH) vehicle was also tested at two different dilutions – the highest concentration used here for drug solubilization was the 1:1000 dilution that did not affect insulin secretion. Data representing the average of all perifusions with the present protocol of untreated controls is included for reference (Avg. Ctrl. All, gray line). (Bottom) Concentration dependence of the effect on insulin secretion as assessed by the effect on the area under the curve of high-glucose induced insulin secretion (AUC_ins_, 11 – 45 min on the top graph), a measure of total insulin secreted. The therapeutic target concentration (*C*_targ_ ≈ 300 nM) and the estimated half-maximal inhibitory concentration (IC_50_ = 10,800 nM) are indicated by red and blue arrows, respectively. Vehicle (here, methanol MeOH) data are shown at the highest CsA concentration where the corresponding dilution was used (e.g., 1:1000 at 25 μM CsA). Data shown are average ± SD (*n* = 3).

**Table 1.**
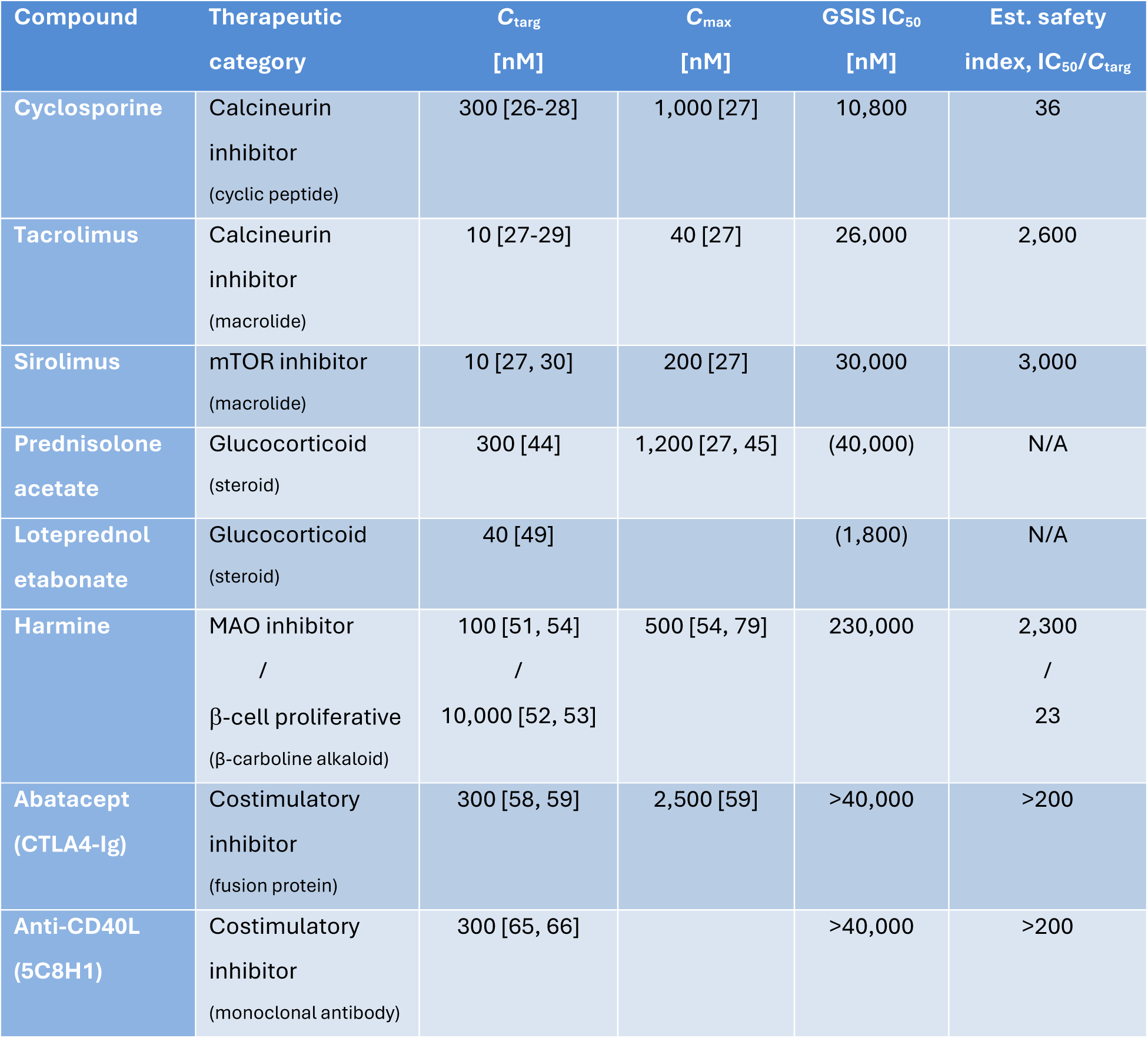
Summary data for the therapeutics included in the present study.

### Calcineurin inhibitors – tacrolimus

Tacrolimus (Prograf®), a macrolide (macrocyclic lactone) CNI, was approved by the FDA in 1994 for use in liver and then kidney transplant. It is more potent than CsA with target blood concentrations in the 4–12 ng/mL (5–15 nM) range [27–29]. It also has a relatively narrow therapeutic index, and blood levels are commonly monitored. Here, we found it to affect islet GSIS in our assay in a manner similar to CsA; however, it became highly detrimental only at concentrations that are considerably higher than its therapeutically effective one. It showed a somewhat strange concentration dependence as it quite consistently caused some (∼10–25%) suppression at the two lowest concentrations tested (40 and 160 nM), but then none at the next two (640 nM and 5.5 μM) leaving even the first-phase response essentially intact. At the two highest concentrations tested here (55 and 109 μM), it essentially wiped out the GSIS response as well as the KCl-induced insulin release indicating that these concentrations are already within the cytotoxic range. Considering that its target blood concentration is *C*_targ_ ≈ 10 nM, about 30-fold less than that of CsA, it did not affect the GSIS profile significantly up to 5 μM (Figure 3, top) and fit of the AUC data suggested an estimated half-maximal inhibitory concentration of IC_50_ = 26,000 nM (Figure 3, bottom). This indicates a ∼2,600-fold separation compared to *C*_targ_, much larger than that obtained for CsA (Table 1).

**Figure 3.**
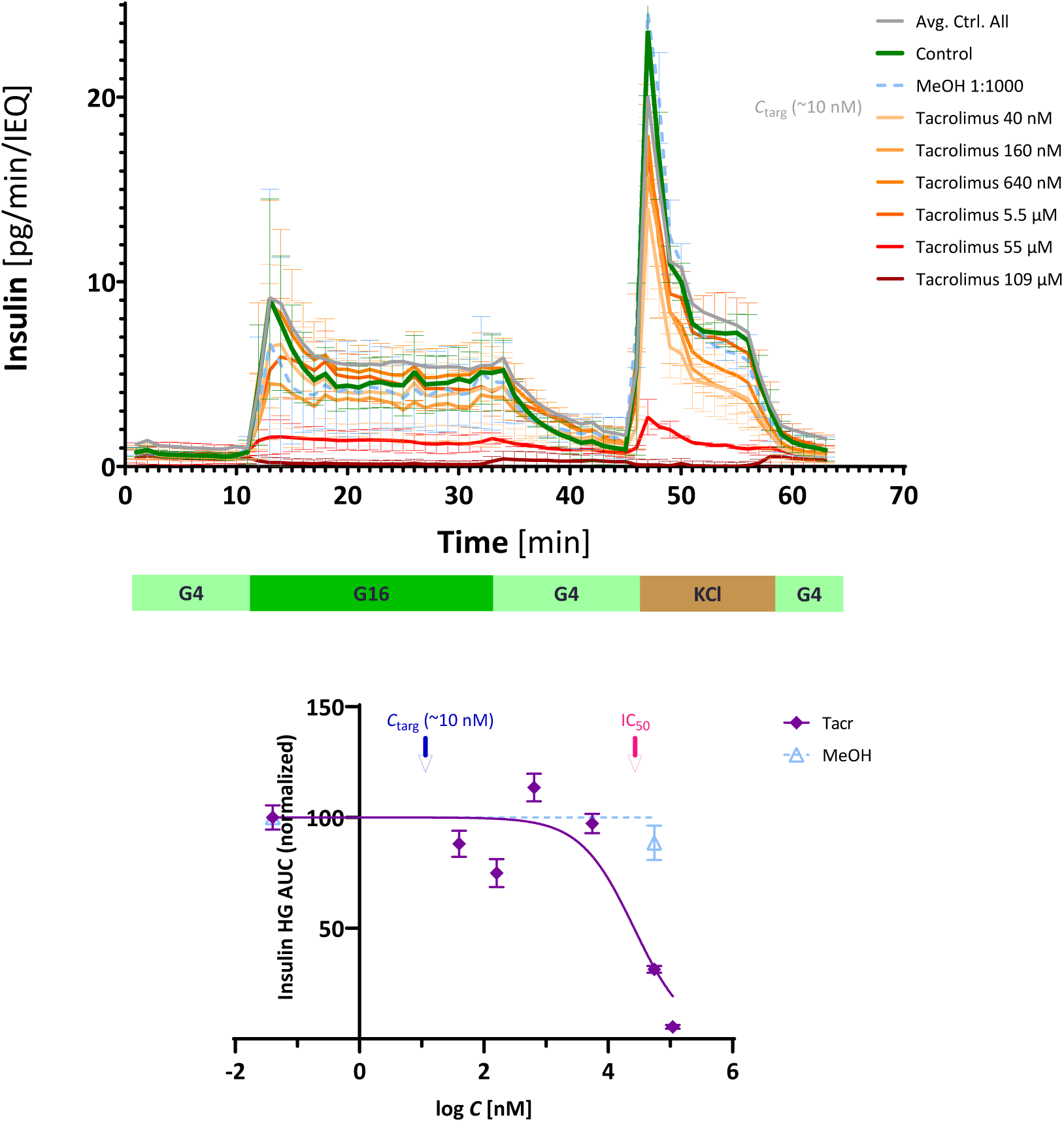
Concentration-dependence of the effect of tacrolimus on the GSIS of human islets. (Top) Time-profile of GSIS of isolated islets exposed to increasing concentrations of tacrolimus for 24 h and then perifused using a stepwise sequence of low-high-low glucose (G4–G16–G4) as indicated. As before, increasing drug concentrations are denoted with increasingly darker colors, and data representing the average of all perifusions with the present protocol of untreated controls is included for reference (Avg. Ctrl. All, gray line). The vehicle (MeOH 1:1000) had no significant effect on insulin secretion. (Bottom) Concentration dependence of the effect on insulin secretion as assessed by the effect on AUC_ins_. The therapeutic target concentration (*C*_targ_ ≈ 10 nM) and the estimated half-maximal inhibitory concentration (IC_50_ = 26,000 nM) are indicated by red and blue arrows, respectively. As before, vehicle (MeOH) data are shown at the highest tacrolimus concentration where the corresponding dilution was used (e.g., 1:1000 at 55 μM). Data shown are average ± SD (*n* = 4).

### mTOR inhibitors – sirolimus

Sirolimus (rapamycin; Rapamune®), another macrolide immunosuppressant with a chemical structure somewhat similar to that of tacrolimus but with a larger ring (31-membered vs 23-membered), is not a calcineurin inhibitor but a mammalian target of rapamycin (mTOR) inhibitor. It was approved by the FDA in 1999 and is indicated for the prevention of organ transplant rejection, prevention of restenosis following balloon angioplasty, and other uses having less nephrotoxicity than CNIs. The target blood concentration range for sirolimus is in the 10–100 ng/mL (10–100 nM) range [27, 30]. Considering such a therapeutic target concentration (*C*_targ_ ≈ 10 nM), it did not affect the GSIS profile at all up to 5,000 nM range (Figure 4, top). Fitting of the AUC data as before suggested an estimated IC_50_ = 30,000 nM (Figure 4, bottom) indicating a ∼3,000-fold separation compared to *C*_targ_, even better than that for tacrolimus. Just as for tacrolimus, the two highest concentrations tested here (55 and 109 μM) essentially wiped out the GSIS response as well as the KCl-induced insulin release indicating that these concentrations are already within the cytotoxic range.

**Figure 4.**
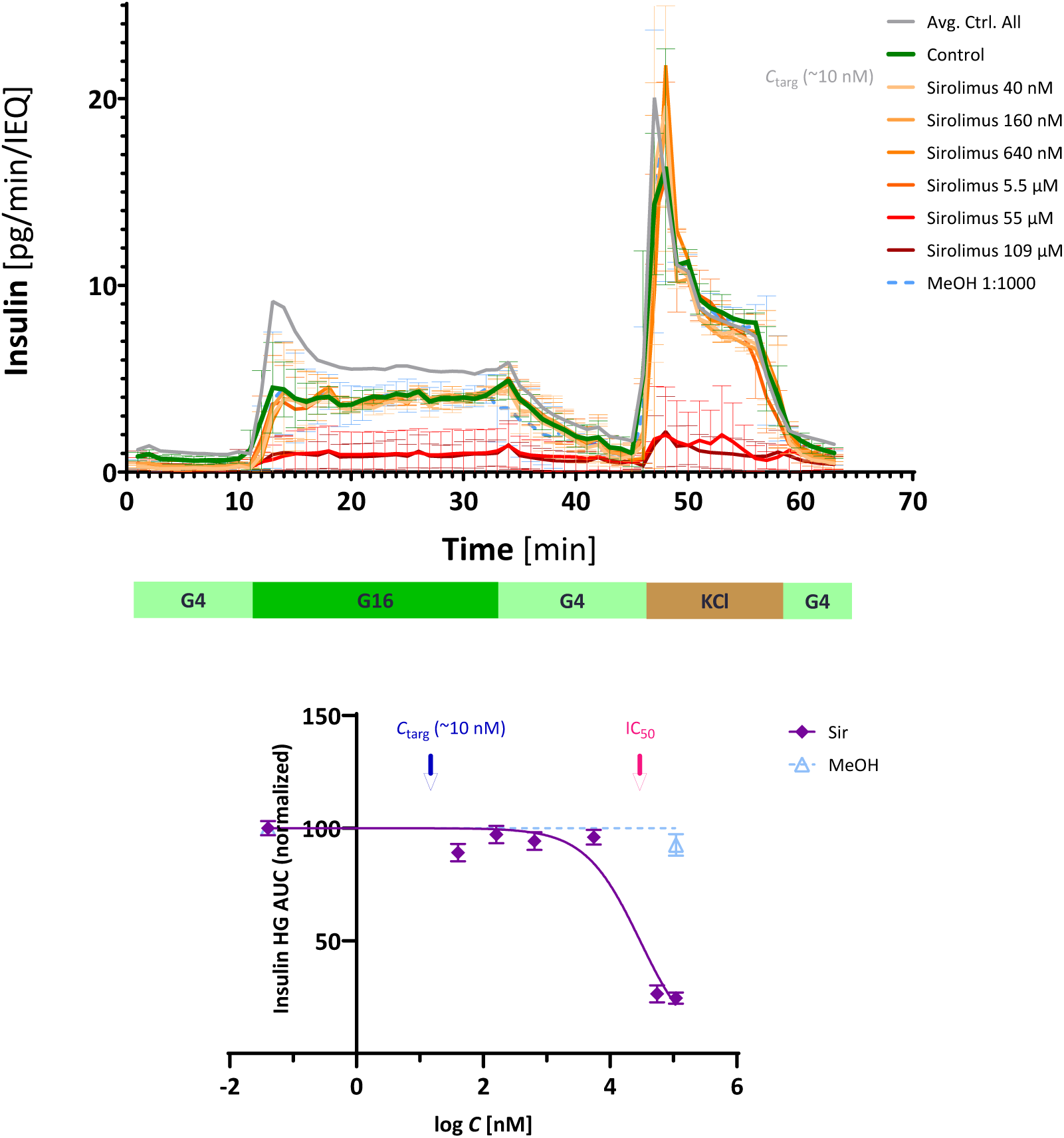
Concentration-dependence of the effect of sirolimus on the GSIS of human islets. (Top) Time-profile of GSIS of isolated islets exposed to increasing concentrations of sirolimus for 24 h and then perifused using a stepwise sequence of low-high-low glucose (G4–G16–G4) as indicated. As before, increasing drug concentrations are denoted with increasingly darker colors. The vehicle (MeOH 1:1000) had no significant effect on insulin secretion. (Bottom) Concentration dependence of the effect on insulin secretion as assessed by the effect on AUC_ins_. The therapeutic target concentration (*C*_targ_ ≈ 10 nM) and the estimated half-maximal inhibitory concentration (IC_50_ = 30,000 nM) are indicated by red and blue arrows, respectively. Vehicle (MeOH) data are shown at the highest sirolimus concentration where the corresponding dilution was used (e.g., 1:1000 at 109 μM). Data shown are average ± SD (*n* = 3).

### Glucocorticoids – prednisolone acetate

Glucocorticoids (GCs), corticosteroids commonly used as immunomodulatory and anti-inflammatory medications, are highly effective therapeutics, but are well known to disturb glucose and lipid homeostasis – side effects that seriously limit their chronic use. A large portion of those treated with GCs long-term and at higher doses (30–40%) develop glucocorticoid-induced diabetes (GCID) [31]. GCs cause hyperglycemia in a dose-dependent manner via several mechanisms that include, among others, the induction or worsening of pre-existing insulin resistance, the increase of hepatic gluconeogenesis, and the stimulation of appetite and weight gain. They have been long known to inhibit insulin release [32–35] as well as to produce whole-body insulin resistance and to exacerbate diabetes following prolonged administration [36]. There is, however, some controversy surrounding GCs, as under certain conditions, they have been found to enhance insulin secretion in islets [37, 38]. There are also claims that GCs have overall beneficial effects on islet transplantation and function by reducing the effects of inflammatory cytokines [39, 40], and there might be a mechanism for local GC regeneration selectively within the β-cells to protect against inflammatory β-cell destruction [41]. Therefore, we felt that there is a particular need for detailed concentration-dependent studies of the effects of GCs on GSIS profile.

Prednisolone acetate (PA), the 21-acetate ester and thus a prodrug of prednisolone, was one of the GCs we evaluated because it is also used as an eyedrop (Pred Forte®), a formulation of particular interest for islet transplantation in the anterior chamber of the eye (ACE), which we have been exploring [42, 43]. The *C*_targ_ of prednisolone is somewhere in the 110 ng/mL (300 nM) range [44, 45]. This is in general agreement with a 10*K*_d_ value (often assumed as needed to ensure full receptor activation as at that concentration ∼90% of all receptors should be occupied [46]) considering its estimated equilibrium dissociation constant *K*_d_ for binding to the GC receptor of 35 nM [47]. These *K*_d_ values have been shown to agree well with the immunosuppressive ability of GCs as measured by whole blood lymphocyte proliferation assay; for example, the IC_50_ of prednisolone for this therapeutically relevant effect was found to be 51 nM, in close agreement with the *K*_d_ = 35 nM value [48].

Compared to the other small molecules assayed here (CNIs and mTOR inhibitors plus harmine, which will be discussed later), the GCs tested here showed a different GSIS inhibitory pattern. They already significantly inhibited GSIS at therapeutic concentrations; however, they did not alter the overall shape of the time-profile (i.e., did not eliminate the first phase or the KCl-induced response) just depressed everything while also never flatlining the entire response even at high concentrations (Figure 5, Figure 6). Considering that the target concentration of PA is *C*_targ_ ≈ 300 nM, it already suppressed somewhat the GSIS profile at 40 nM (by ∼15%) and by almost 50% at 10.2 μM, the highest concentration tested here (Figure 5, top). Contrary to the previous cases, the insulin AUC data cannot be fitted well with a classic law of mass action type sigmoid function (i.e., *n*_Hill_ = 1) as it only shows a slow increase of effect with increasing concentrations. Fit as shown here (Figure 5, bottom) was obtained with *n*_Hill_ = 0.25 indicating a very slow concentration-dependency. Hence, the large and ill-defined IC_50_ estimate obtained (40,000 nM = 40 μM), which should not be considered a reliable quantitative estimate (e.g., the 95% confidence interval, CI, is 15,000 to 116,000 nM).

**Figure 5.**
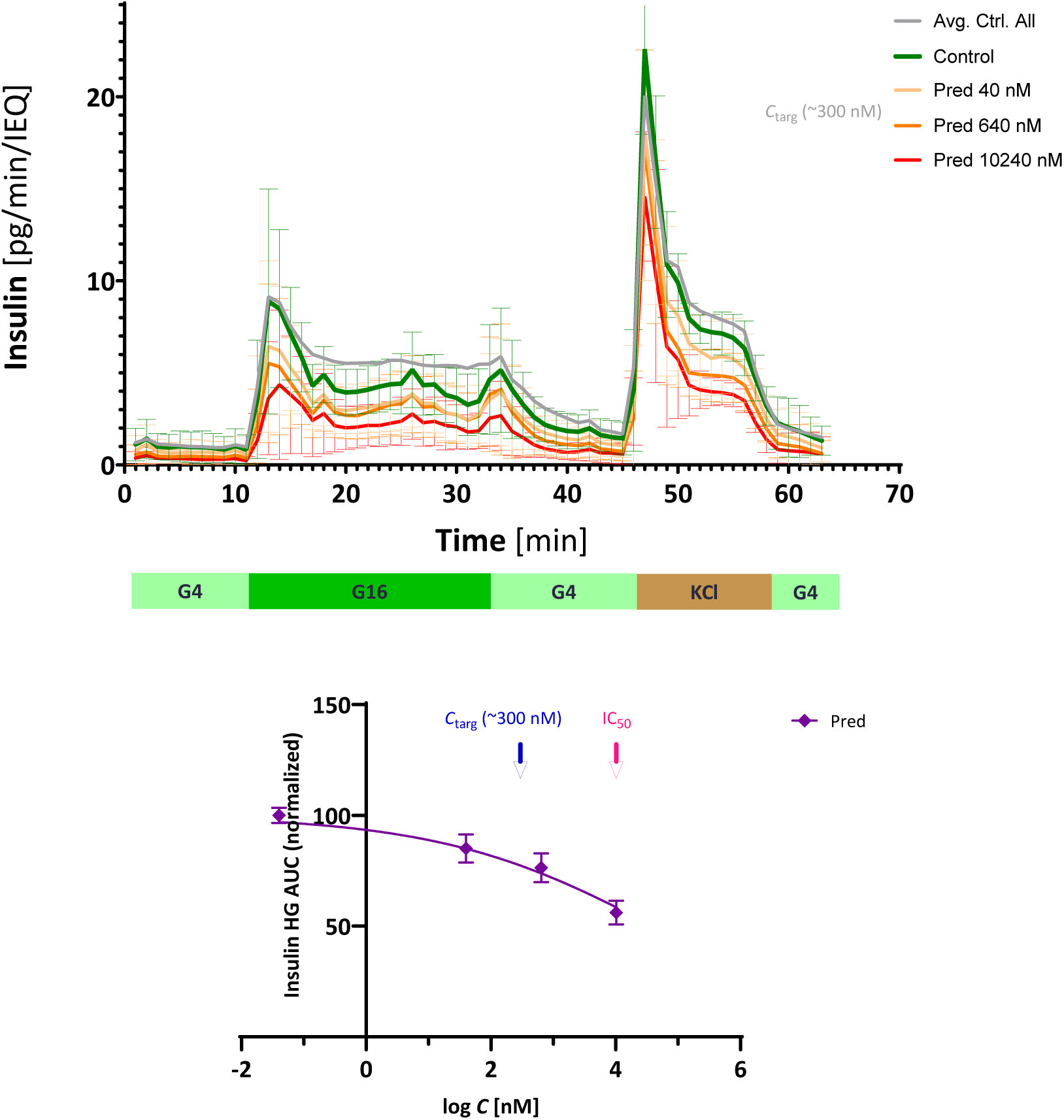
Concentration-dependence of the effect of prednisolone acetate on the GSIS of human islets. (Top) Time-profile of GSIS of isolated islets exposed to increasing concentrations of prednisolone acetate (Pred) for 24 h and then perifused using a stepwise sequence of low-high-low glucose (G4–G16–G4) as indicated. As before, increasing drug concentrations are denoted with increasingly darker colors. (Bottom) Concentration dependence of the effect on insulin secretion as assessed by the effect on AUC_ins_. The therapeutic target concentration (*C*_targ_ ≈ 300 nM) and the estimated half-maximal inhibitory concentration (IC_50_ = 40,000 nM) are indicated by red and blue arrows, respectively. Fit shown here for the AUC data was obtained with *n*_Hill_ = 0.25. Data shown are average ± SD (*n* = 8).

**Figure 6.**
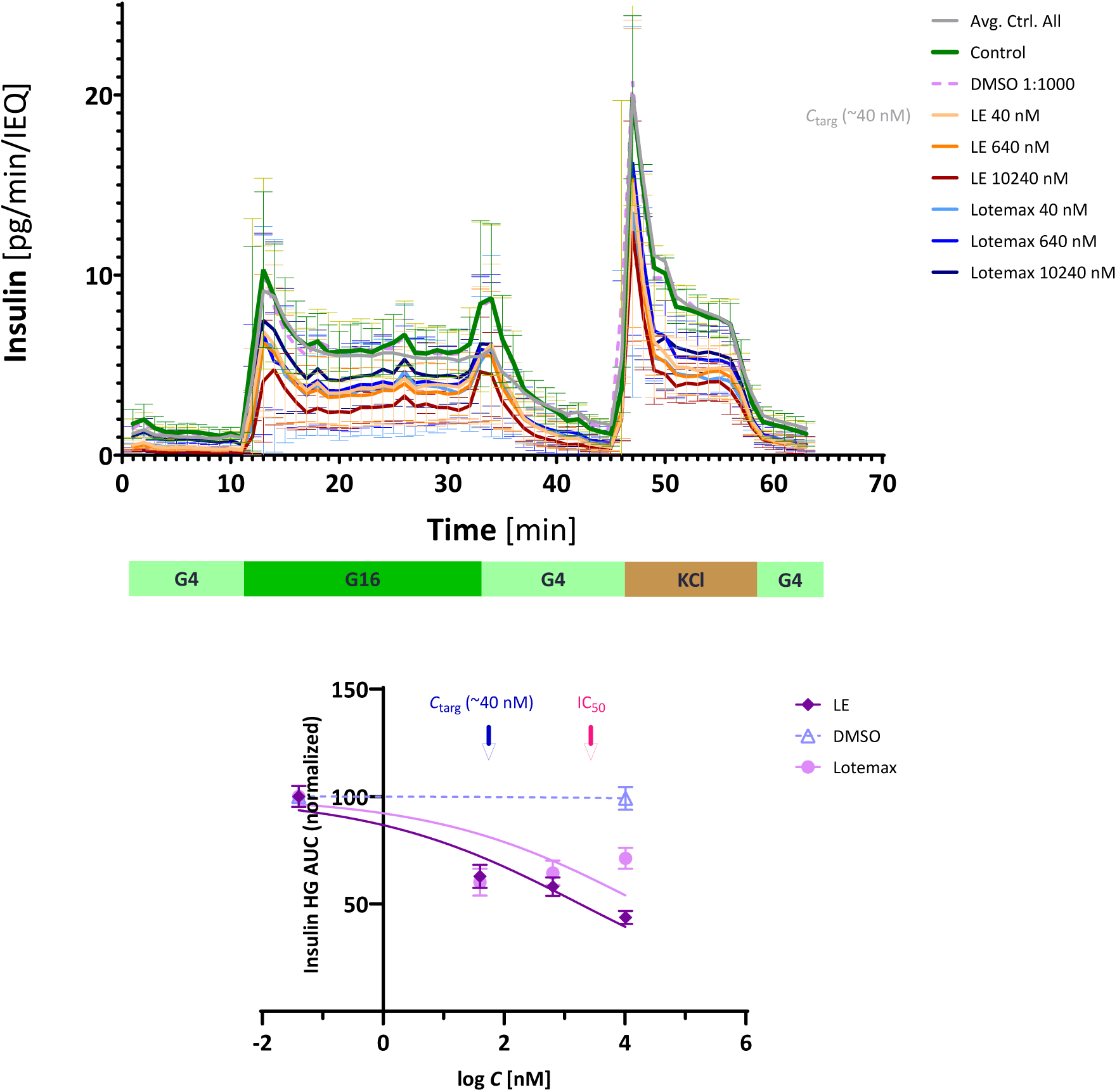
Concentration-dependence of the effect of loteprednol etabonate on the GSIS of human islets. (Top) Time-profile of GSIS of isolated islets exposed to increasing concentrations of loteprednol etabonate either as its research or clinically approved formulation (LE and Lotemax, respectively) for 24 h and then perifused using a stepwise sequence of low-high-low glucose (G4–G16–G4) as indicated. As before, increasing drug concentrations are denoted with increasingly darker colors. The vehicle (DMSO 1:1000) had no significant effect on insulin secretion. (Bottom) Concentration dependence of the effect on insulin secretion as assessed by the effect on AUC_ins_. The therapeutic target concentration (*C*_targ_ ≈ 40 nM) and the estimated half-maximal inhibitory concentration (IC_50_ = 1,800 nM) are indicated by red and blue arrows, respectively. Fit shown here for the AUC data was obtained with *n*_Hill_ = 0.25. Vehicle (here, DMSO) data are shown at the concentration where the corresponding dilution was used (e.g., 1:1000 at 10.24 μM). Data shown are average ± SD (*n* = 4).

### Glucocorticoids – loteprednol etabonate

The other GC explored in this study was loteprednol etabonate (LE), which is also approved for clinical use as an eye drop (Lotemax®). LE is a soft drug developed to have reduced side effects compared to other GCs following topical administration [49, 50]; thus, it is in some ways the conceptual opposite of PA, which is a prodrug: while prodrugs are inactive as administered and need to be metabolically activated, soft drugs are active and designed to be rapidly inactivated after exerting their intended therapeutic action [50]. Since LE is only applied locally, it does not have an accepted systemic therapeutic target blood concentration, but considering its estimated equilibrium dissociation constant *K*_d_ of 4 nM [47] and the same calculation as for PA (∼10*K*_d_), 40 nM (20 ng/mL) can serve as a reasonable guideline. Just as with PA, LE also did not alter the overall shape of the GSIS time-profile just depressed it, and it also did not flatline the entire response even at high concentrations (10.2 μM, Figure 6). At *C*_targ_ level, LE suppressed the GSIS profile by ∼40%, and at the highest concentration tested (10.2 μM) by 60% (Figure 6). Fitting of the AUC data suggested an estimated IC_50_ = 1,800 nM, but, again, this cannot be considered a reliable estimate as it resulted from a very inadequate fit (e.g., 95% CI of 190 to 20,000 nM) – even more so than in the case of PA (Figure 6, bottom). It was obtained with *n*_Hill_ = 0.25, the same value that was used for PA. For LE, in addition to its research only formulation (acquired from Sigma-Aldrich), we also tested its clinically available formulation (Lotemax) at the same concentrations. While this gave very similar results for the two lowest concentrations (40 and 640 nM), it resulted in less suppression at the highest concentration (10.2 μM) (Figure 6).

### Harmine

In this study, we also investigated harmine, a tricyclic alkaloid (i.e., naturally occurring nitrogen-containing compound) that has several biological activities of interest including antitumor, neuroprotective, antiparasitic, anti-inflammatory, and antidiabetic properties [51] including its potential ability to induce β-cell proliferation and increase islet mass [52, 53]. Harmine inhibits monoamine oxidase (MAO) with an IC_50_ of ∼5 nM and DYRK1A with an IC_50_ of ∼100 nM [54]; for β-cell proliferation, it is suggested to be used at the considerably higher concentration of 10 μM (2,100 ng/mL) [52, 53]. Here, we explored its effect on dynamic GSIS at concentrations ranging from 2 to 500 μM (Figure 7). As noticeable in the figure, concentrations of up to 50 μM did not affect the GSIS profile, but higher concentrations (250 and 500 μM) seriously depressed it, eliminating the first-phase response and also diminishing the KCl-induced insulin release (Figure 7, top). Fitting of the AUC data gave an IC_50_ = 230 μM (Figure 7, bottom) – a safe separation from the MAO and DYRK1A IC_50_s, but only a 23-fold separation compared to the *C*_targ_ for β-cell proliferation.

**Figure 7.**
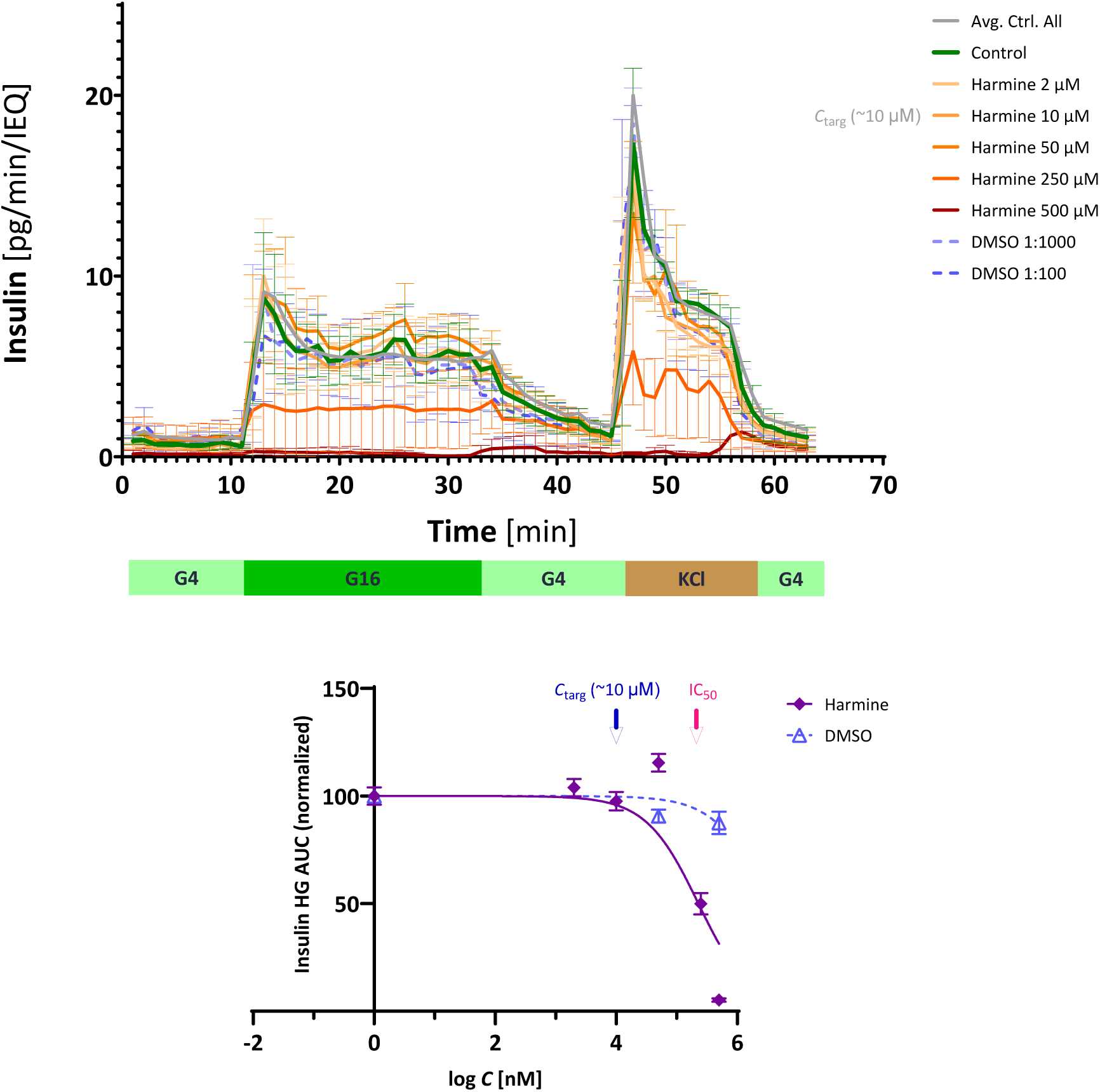
Concentration-dependence of the effect of harmine on the GSIS of human islets. (Top) Time-profile of GSIS of isolated islets exposed to increasing concentrations of harmine for 24 h and then perifused using a stepwise sequence of low-high-low glucose (G4–G16–G4) as indicated. As before, increasing drug concentrations are denoted with increasingly darker colors. The vehicle (DMSO 1:1000 and 1:100) had no significant effect on insulin secretion. (Bottom) Concentration dependence of the effect on insulin secretion as assessed by the effect on AUC_ins_. The therapeutic target concentration (*C*_targ_ ≈ 10 μM) and the estimated half-maximal inhibitory concentration (IC_50_ = 230 μM) are indicated by red and blue arrows, respectively. Vehicle (DMSO) data are shown at the concentrations where the corresponding dilutions were used. Data shown are average ± SD (*n* = 3).

### Biologics – abatacept (CTLA4-Ig)

In addition to classic small-molecule drugs, several biologics are also routinely used for immunosuppressive therapies as the number of FDA approved antibodies and other biologics has been steadily increasing since the 1990s [55]. The use of biologics in general is often hindered by issues such as immunogenicity [56] and that of immunomodulatory biologics in particular by the high likelihood of unwanted adverse effects, such as cytokine release syndrome, serious / protracted infections, malignancy, or anaphylaxis [57]. However, they are much less likely to be toxic to β-cells and affect insulin secretion. Abatacept (Orencia®) is a fusion protein composed of the Fc region of immunoglobulin G1 (IgG1) fused to the extracellular domain of human cytotoxic T-lymphocyte-associated antigen 4 (CTLA-4, CD152) that binds to CD80/86 inhibiting the CD80/86–CD28 costimulatory interaction needed for T cell activation [58]. It received FDA approval in 2005 and is used for RA treatment. Target serum concentrations are around 30,000 ng/mL (330 nM) [58, 59]. Here, we tested it for concentrations ranging from 40 nM to 5,000 nM and found that it had no noticeable effect on the GSIS profile of human islets (Figure 8). Thus, we could not obtain a reliable IC_50_ estimate, but it is certain that IC_50_ > 40,000 nM (Figure 8, bottom), several orders of magnitude larger than *C*_targ_ (∼300 nM), indicating that inhibition of insulin secretion is unlikely to be a concern for abatacept treatment.

**Figure 8.**
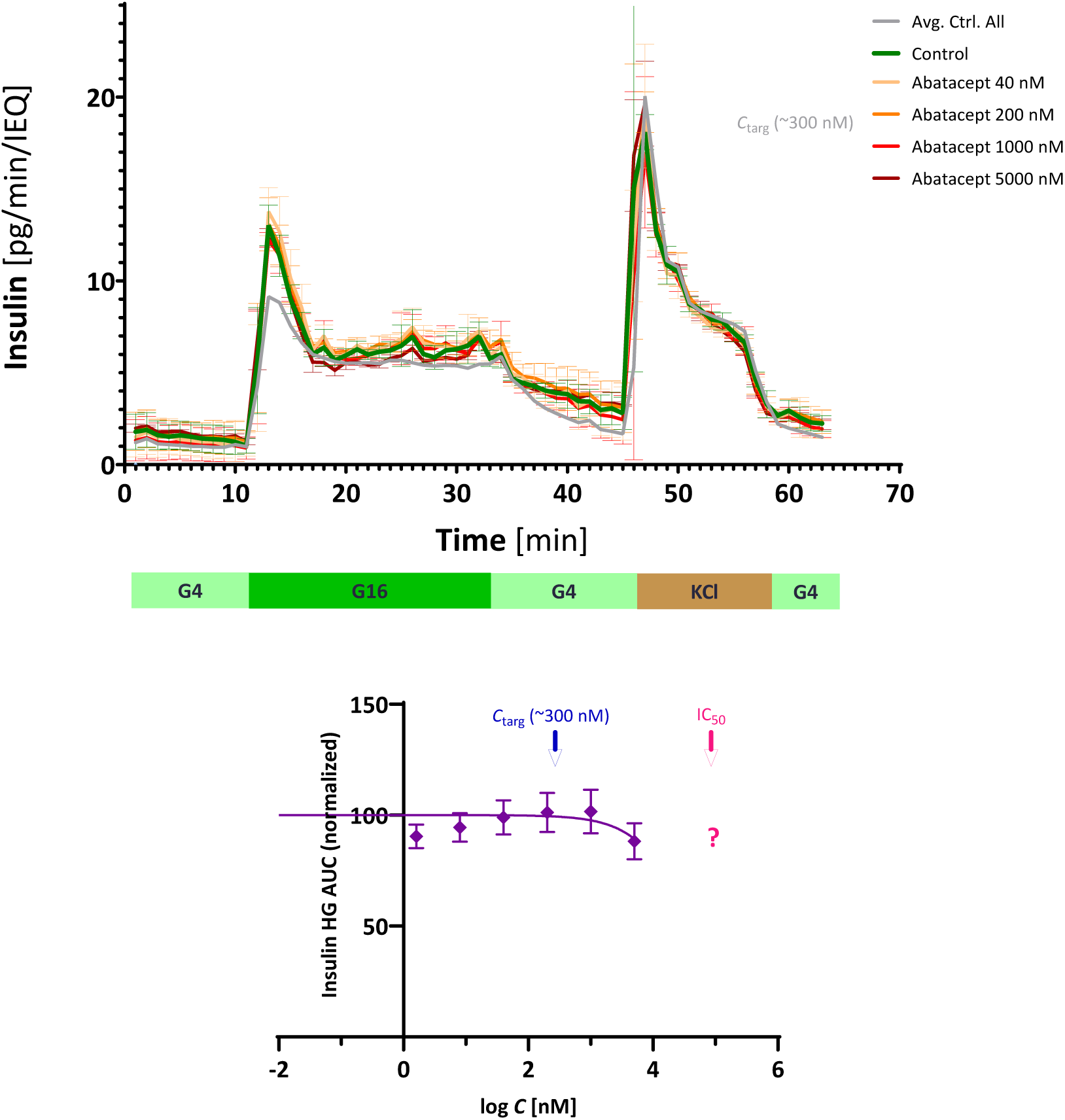
Concentration-dependence of the effect of abatacept (CTLA4-Ig) on the GSIS of human islets. (Top) Time-profile of GSIS of isolated islets exposed to increasing concentrations of abatacept for 24 h and then perifused using a stepwise sequence of low-high-low glucose (G4–G16–G4) as indicated. As before, increasing drug concentrations are denoted with increasingly darker colors. (Bottom) Concentration dependence of the effect on insulin secretion as assessed by the effect on AUC_ins_. The therapeutic target concentration (*C*_targ_ ≈ 300 nM) and a rough estimate of the half-maximal inhibitory concentration (IC_50_ > 40,000 nM) are indicated by red and blue arrows, respectively. Data shown are average ± SD (*n* = 3).

### Biologics – anti-CD40L

Inhibition of the CD40–CD40L costimulatory interaction is also of interest as an immunomodulatory therapy in general [60] and for pancreatic islet transplantation in particular, as it has been shown in multiple animal models to allow the engraftment of allogeneic (and sometimes even xenogeneic) islets resulting in long-term insulin independence with the possibility of induced operational tolerance [10]. The first-generation of CD40L antibodies such as ruplizumab (hu5c8) were abandoned due to their thrombolytic side effects, but with the recognition that this was driven by their Fc region and can be avoided, there is a strong resurgence of interest [61] and several second-generation Fc-silent anti-CD40L antibodies are in advanced clinical trials. They include, among others, letolizumab, dapirolizumab pegol, frexalimab, tegoprubart, and TNX-1500 (Tonix). Tegoprubart is in Phase 2a trials for amyotrophic lateral sclerosis (ALS) [62, 63] and Phase 1/2a trial for β-cell replacement with very promising results [64]. There is no clear target serum concentration as it is not approved for clinical use, but 300 nM can serve as a reasonable estimate [65, 66].

Here, we tested the effect of anti-CD40L (5C8H1) at concentrations ranging from 1.6 nM to 5,000 nM and found that, similar to abatacept, it had no effect on the GSIS profile of human islets (Figure 9). Therefore, just as for abatacept, we could not obtain a reliable IC_50_ estimate, but it is certain that IC_50_ > 40,000 nM (Figure 9, bottom) so that inhibition of insulin secretion is not a concern for anti-CD40L antibody either.

**Figure 9.**
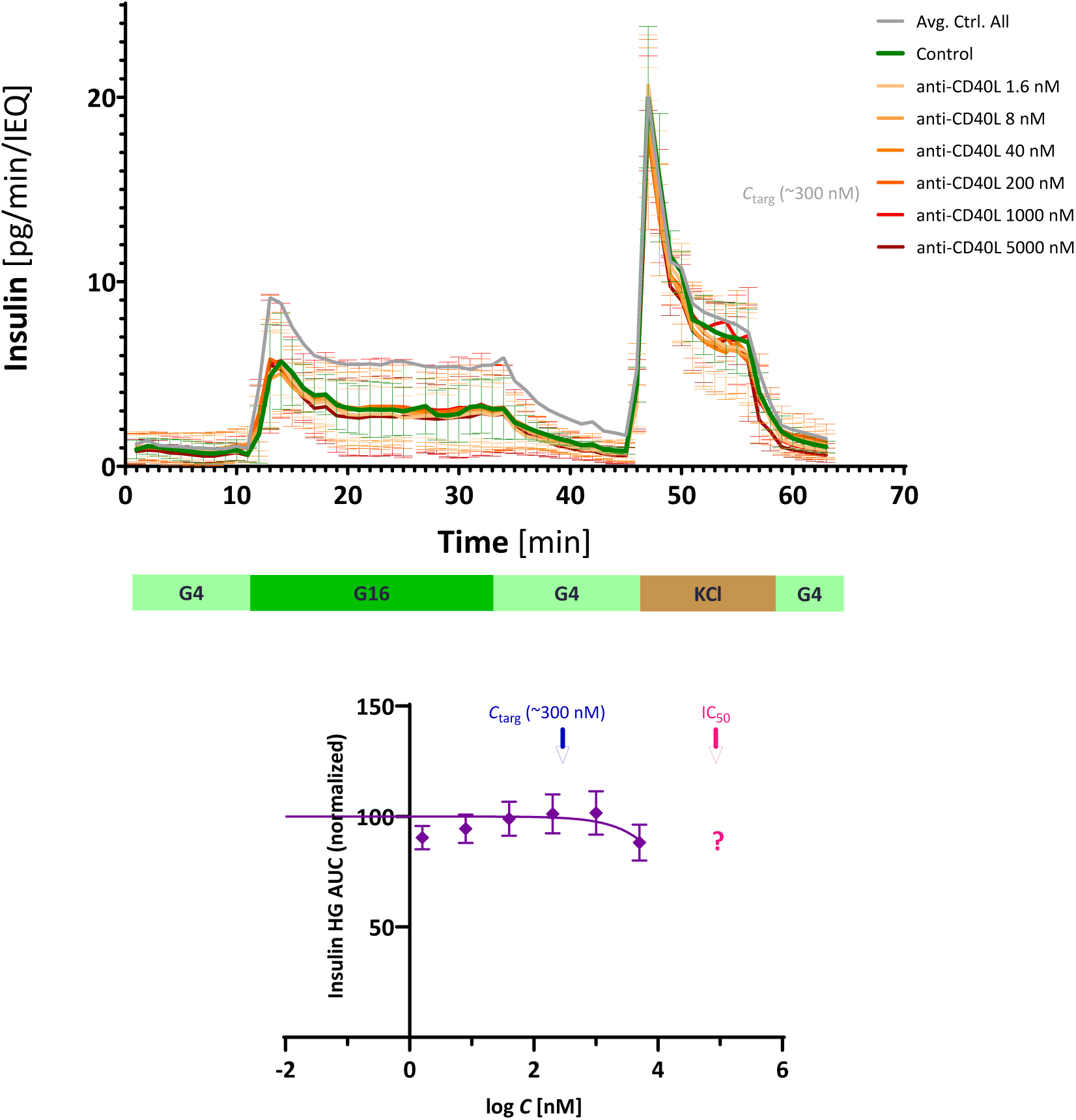
Concentration-dependence of the effect of anti-CD40L antibody (5C8H1) on the GSIS of human islets. (Top) Time-profile of GSIS of isolated islets exposed to increasing concentrations of anti-CD40L antibody for 24 h and then perifused using a stepwise sequence of low-high-low glucose (G4–G16–G4) as indicated. As before, increasing drug concentrations are denoted with increasingly darker colors. (Bottom) Concentration dependence of the effect on insulin secretion as assessed by the effect on AUC_ins_. The therapeutic target concentration (*C*_targ_ ≈ 300 nM) and a rough estimate of the half-maximal inhibitory concentration (IC_50_ > 40,000 nM) are indicated by red and blue arrows, respectively. Data shown are average ± SD (*n* = 3).

## Discussion

Considering the well-known propensity of immunosuppressive therapies to show β-cell toxicity, inhibit insulin secretion, and induce PTDM, it is surprising that only a very limited number of studies have been done to characterize the concentration-dependence of these effects – an essential part of pharmacological / toxicological characterization. Furthermore, many of these studies have been done with rodent and not human islets while there are significant differences in their behavior [16]. Here, to characterize in detail the concentration-dependent suppression of GSIS, we performed dynamic perifusion experiments with isolated human islets incubated with increasing concentrations of several important clinically used immunosuppressive drugs that included small-molecule drugs such as cyclosporine, sirolimus, tacrolimus, prednisolone acetate, and loteprednol etabonate, as well as biologics, namely CTLA4-Ig (abatacept) and anti-CD40L.

Human islets show large, several-fold variability in their insulin secretion time profiles as assessed in dynamic perifusion assay – this is evident from the large error bars of Figure 1 and was illustrated in more detail, for example, by the spaghetti plots of individual profiles in our earlier paper (*n* = 70 untreated control islets, Supplementary Figure S6) [17] or the detailed data of an even larger sample (*n* = 299) assessed by the IIDP [25]. Despite this and despite differences in the standard protocols used, overall secretion profiles of untreated islets are quite similar as shown in Figure 1. The main difference between the profiles obtained by us and IIDP is likely due to islets in our experiments being from a more stringent selection criteria regarding viability, purity, and donor BMI and being less stressed due to the different equipment and protocol used resulting in somewhat lower baseline and stimulated insulin secretions as well as less variability (lower standard deviations).

### Biologics

We found that the biologics tested here (abatacept and anti-CD40L) showed no significant detrimental effects on the insulin secreting ability of human islets after 24 h of treatment even at relatively high concentrations (5 μM) (Figure 8, Figure 9). Accordingly, GSIS inhibitory IC_50_ values could not be reliably established from these data, and the values obtained here (>40 μM; Table 1) can be considered only as rough estimates. In agreement with these observations, a recent study on localized immunomodulation using a subcutaneous vascularized device (NICHE) that included perifusion studies also found no significant effects on GSIS for abatacept and anti-CD40L (MR-1) at concentrations of 0.1, 0.5, and 1.0 mg/mL (so up to 11,000 nM for abatacept and 7,000 nM for anti-CD40L) after 5 days of incubation with rodent islets [67]. The same study also evaluated three other biologics (anti-lymphocyte serum ALS, anti-CD2, and anti-IL6) and found no significant effect on GSIS for them either for similarly high concentrations even after 5 days of incubation [67]. Therefore, while immunogenicity and other serious side effects seen with immunomodulatory biologics remain an important concern [56, 57], inhibition of insulin secretion is unlikely to be a major concern for most highly specific biologic treatments.

### Calcineurin and mTOR inhibitors

On the other hand, all small-molecule drugs tested here inhibited insulin secretion in a concentration-dependent manner. The evaluated calcineurin and mTOR inhibitors (cyclosporine, tacrolimus, and sirolimus) did not affect GSIS at concentrations in their therapeutic range but completely inhibited it at higher concentrations. Their inhibitory effects on the total insulin secreted (evaluated as the AUC data from the corresponding perifusion time-profiles) followed classic law of mass action type responses, i.e., could be fitted well with sigmoid curves that have a unity Hill slope 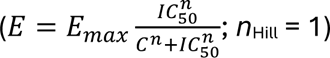 on the semi-log scale; these were used to estimate the half-maximal inhibitory concentration values, IC_50_s, obtained here (Figure 2, Figure 3, Figure 4; bottom). Among the immunosuppressive drugs tested here, CsA showed the least separation between its therapeutic target (*C*_targ_) and median inhibitory (IC_50_) concentrations. We found that concentrations not much higher than *C*_targ_ (300 nM) already affected insulin secretion as first-phase secretion was blunted and total insulin secretion was diminished at 5,000 nM (5 μM; Figure 2, top) and the IC_50_ for total insulin secretion (AUC) was estimated to be around 10,000 nM. Thus, there was an only ∼35-fold separation between the desired *C*_targ_ and the GSIS inhibitory IC_50_, far less than for tacrolimus and sirolimus (Table 1). In agreement with our results, a study by Bugliani and co-workers at the University of Pisa found that CsA at 150 ng/mL (125 nM; 96 h incubation), a concentration within the therapeutic range, did not inhibit GSIS of human islets [68]. On the other hand, a perifusion study with canine islets at the University of Texas found that CsA at 100 nM (72 h) already significantly inhibited both first- and second-phase insulin release [69].

Tacrolimus and sirolimus showed highly detrimental effects, but only at concentrations much higher than their therapeutic target levels (*C*_targ_ << IC_50_). Their estimated GSIS IC_50_s obtained here are >1,000-fold higher than their commonly accepted target levels (*C*_targ_), indicating that they do not significantly inhibit insulin secreting ability, probably up to their cytotoxic concentrations. Tacrolimus consistently seemed to show some possible GSIS inhibition at the two lowest concentrations tested (40 and 160 nM), which are in the therapeutic range, but the next two highest (640 nM and 5.5 μM) did not show any inhibition, leaving even the first-phase response essentially intact (Figure 3). Hence, we obtained a quite high overall estimate for IC_50_ (26,000 nM; Figure 3, Table 1). In general agreement with our results, a few other previous studies found relatively low concentrations of tacrolimus to inhibit GSIS. For example, the previously mentioned University of Pisa study found that tacrolimus at 10 ng/mL (13 nM; 96 h) significantly inhibited GSIS of human islets [68]. Another study from Oslo University found that it significantly inhibited static GSIS of human islets at 30 ng/mL (37 nM; 24 h) worsened by addition of a glucocorticoid (methylprednisolone, 1,000 ng/mL) [70]. A study from UCLA also found that it inhibited GSIS of human islets at 50 and 100 ng/mL (∼60 and 120 nM; 48 h) [71], while a perifusion study at Kyoto University with rat islets suggested some inhibition already at 3 nM (24 h) [72]. A perifusion study at the University of British Columbia found significant inhibition on perifusion GSIS at 37 nM (24 h) with human islets as well as at 10,000 nM (acute) with mouse islets [73]. On the other hand, the perifusion study with canine islets at the University of Texas mentioned earlier found that contrary to CsA, tacrolimus at 100 nM (72 h) showed only little inhibition of insulin release [69].

While sirolimus negatively affected islet function *in vivo* in some studies [74, 75], it did not in others [76]. Here, we found no detrimental effect up to quite high concentrations (Figure 4). In agreement with this, the mentioned perifusion study at University of British Columbia found no significant effects for sirolimus on perifusion GSIS at 33 nM (24 h) with human islets as well as at 1,000 nM (acute) with mouse islets [73]. Among previous studies of its effect on GSIS, the Oslo University study mentioned earlier also found that sirolimus showed some but not significant inhibition of static GSIS at 30 ng/mL (33 nM; 24 h); this, however, was worsened by addition of a glucocorticoid (methylprednisolone, 1,000 ng/mL)[70]. In the UCLA study, sirolimus at 1 ng/mL did not affect insulin secretion, but at 50 ng/mL (55 nM; 48 h) it did significantly inhibit both the baseline and the high-glucose induced insulin secretion of human islets [71].

### Glucocorticoids

GCs showed a distinctly different pattern as they did not distort the shape of the time-profile but inhibited overall GSIS already at therapeutic levels depressing both the first- and second-phase as well as KCl-induced insulin secretion to the same degree. We found that for the glucocorticoids tested here (PA and LE), the suppression was already present even at concentrations that are in their therapeutic range, but it was a general suppression of the entire time-profile without altering the shape. Whereas, the previously discussed immunosuppressive agents did not affect GSIS at therapeutic concentrations and then clearly changed the shape of the response suppressing first the first-phase response at higher concentrations and entirely wiping out the response at even higher concentrations, GCs showed a different pattern (e.g., Figure 2 or Figure 4 vs Figure 5 or Figure 6). While there was a concentration-dependent tendency in the suppressive effects of both glucocorticoids tested here, the responses did not fit a classic law of mass action type sigmoid response (*n*_Hill_ = 1) as those before: they showed a much slower increase with concentration than expected; hence, indicating *n*_Hill_ < 1 if the same functional form, 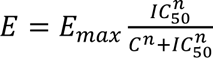, is used (e.g., Figure 5 and Figure 6 versus Figure 2 and Figure 4). For fitting of the AUC data, we used *n*_Hill_ = 0.25 that gave adequate fit for both PA and LE – this corresponds to a very slow concentration-dependency; hence, the large and ill-defined (i.e., wide 95% CIs) IC_50_ estimates obtained. At concentrations corresponding to their target therapeutic levels (*C*_targ_), these GCs already suppressed insulin secretion by 25–40% (Figure 5, Figure 6). In general agreement with this, a recent perifusion study with human islets by Tijani and coworkers from the University of Lille, France gave somewhat similar results as they found prednisolone to suppress GSIS by about 50% at doses of 250, 500, and 1,000 nM (24 h), but without showing a concentration-dependent increase in effect [35]. In the same study, hydrocortisone (HC; cortisol) and dexamethasone (DEX) also caused similar effects when used at therapeutically relevant concentrations.

GCs exert multiple activities, but because their main actions are mediated via the same receptor, which is ubiquitously distributed, both therapeutic as well as unwanted side effects tend to run in parallel and are closely related to potency as measured by receptor-binding affinity (RBA) so that effects at doses adjusted for their relative potency (rRBA) are very similar [47]. For example, compared to DEX, prednisolone is 5–7-times less potent, HC is 10–25 times less potent, while LE is about equipotent [47]. Several *in vitro* studies have shown that GC exposure affects β-cell function and inhibits GSIS in time- and dose-dependent manner (see references in [35]). As mentioned, Tijani and coworkers found depression of the secretion profile (both phases) of human islets at therapeutically relevant GC concentrations (e.g, 250–1,000 nM, 24 h) – similar to our observations but without a concentration-dependent increase in effect [35]. A study with mouse islets by Henquin and coworkers at the University of Bruxelles suggested an IC_50_ of around 20 nM for DEX (18 h treatment), which is about equipotent with LE, with a maximum of 80% inhibition at 250 nM [32]. They found insulin content to not be affected by GC treatment; in fact, to be actually higher in the DEX-treated islets. Perifusion in the same study found a smaller, about 50% suppression of the secretion profile (both phases, thus without altering the shape) after treatment with 1,000 nM DEX (18 h) in general agreement with our data obtained with human islets. Ultimately, they suggested that GCs exert direct genomic effects on β-cells resulting in a generalized inhibition of insulin secretion that is brought about by a decrease in the effectiveness of cytoplasmic calcium on the secretory process [32]. Another perifusion study with rat islets at Yale also found similar strong depression of both phases (64% and 74% for first- and second phase, respectively) caused by 1,000 nM DEX (3 h) without affecting insulin content as well as depression caused by KCl (30 mM) [34]. They also concluded that GC pretreatment impairs insulin secretion via a genomic action, but a clear mechanism was not identified.

### Harmine

Harmine and harmaline are β-carboline alkaloids that occur in different plants (especially in *Peganum harmala* and *Banisteriopsis caapi*) and are part of ayahuasca, a hallucinogenic herbal preparation used for therapeutic and divination purposes by indigenous Amazonian tribes. They are MAO-A inhibitors that have psychotropic properties; thus, can cause dose-related side effects [77]. Harmine is being investigated for several possible therapeutic activities of interest [51]; we included it here due its potential ability to induce β-cell proliferation and increase islet mass [52, 53]. As the β-cell regenerative effect requires relatively high concentrations (*C* > 10,000 nM), harmine presents a critical balance between therapeutic proliferation and functional preservation. With an estimated IC_50_ of 230 μM, harmine exhibits a relatively narrow 23-fold safety index compared to its suggested proliferation target (Table 1). Consequently, for harmine to be effectively utilized in β-cell replacement strategies or localized delivery therapies, dosing will have to be meticulously calibrated. A Phase 1 clinical trial found oral doses up to 2.7 mg/kg safe with those higher associated with adverse effects including vomiting, drowsiness, and limited psycho-activity [78]. Note that another recent study found that peak harmine blood levels after a similar dose (180 mg) administered via a transmucosal delivery system only reached 49 ng/mL (250 nM) in healthy volunteers [79], indicating that other side effects might be more limiting than those on insulin secretion. Harmine blood levels found in traditional ayahuasca users were in the 500 nM range [54], still well below the 10,000 nM suggested for possible β-cell proliferation [52, 53]. Hence, achieving proliferative *C*_targ_ levels will likely require pancreas-targeted or -localized delivery to avoid the otherwise limiting systemic adverse effects.

### Summary and limitations

As the side effects of the required chronic immunosuppression are one of the major factors limiting the applicability of organ and cell transplantation, it is of critical relevance to characterize their dose- and concentration-dependence to optimize therapeutic regimens and clinical protocols. PTDM and inhibition of insulin secretion are important unwanted side effects [2–4] – here, we described a detailed quantification of the concentration dependency of the latter using dynamic perifusion studies with human islets. We found that the inhibitory effect of calcineurin and mTOR small-molecule inhibitors follows a classic sigmoid law of mass action pattern (*n*_Hill_ = 1) with no effect at their therapeutic level that then increases progressively and can be characterized well by half-maximal inhibitory concentration values (IC_50_) as shown by the effect on total insulin secretion (AUC_ins_), for example, in Figure 2 and Figure 4. Corticosteroids showed a different pattern: they already inhibited the amount of insulin secreted even in their therapeutic range but did not alter the overall shape of the time-profile, just suppressed it. Also, their inhibitory effect increased only very slowly with concentration suggesting *n*_Hill_ << 1.

These results are relevant for immunosuppressive therapy following transplantation in general, but particularly for β-cell replacement therapies, which have been shown to restore metabolic control more efficiently than exogenous insulin administration and prevent complications associated with long-term T1D [5–7] but are limited by the negative effects on the engraftment, function, and survival of transplanted islets of all existing systemic immunosuppression regimens [8, 9]. Localized immunosuppression is challenging, but following multiple initial failures, increasing success has been achieved more recently [80–82]; nevertheless, localized delivery still exposes the transplanted cells to high drug concentrations so that GSIS inhibitory effects such as those characterized here remain a major concern that has to be addressed.

Among the limitations of our study, it has to be mentioned that these are *in vitro* experiments utilizing isolated islets removed from their natural systemic vascular and nervous networks; hence, even though islets function well as mini organs on their own, they do not fully reproduce all aspects of the full *in vivo* system. Also, the 24 h drug incubation period used here, while sufficient for observing acute and sub-acute toxicity, may not fully replicate the effects of chronic exposure in patients requiring lifelong immunosuppression.

## Conclusion

The present work demonstrates that targeted biologics preserve human islet glucose-stimulated insulin secretion (GSIS), whereas small-molecule immune suppressants and the possible regenerative agent harmine exhibit distinct, concentration-dependent inhibition. Calcineurin and mTOR inhibitors did not significantly affect GSIS at therapeutic concentration levels (∼*C*_targ_) but increasingly inhibited it at higher concentrations following a typical sigmoid pattern (i.e., well-described by IC_50_ values). Glucocorticoids globally suppressed GSIS even at therapeutic levels and their effect increased only slowly with concentrations. Our findings underscore the critical need for precisely calibrated dosing to safely balance immune protection, tissue regeneration, graft function, and insulin secreting ability in clinical therapies.

## Author Contributions

PB originated and designed the project, conceived the study, designed experiments, provided study guidance, analyzed and interpreted data, and wrote the manuscript; STC, BW, and OA conducted experiments, collected data, and revised and proofread the manuscript.

## Acknowledgments

The authors gratefully acknowledge the organ donors and their families for their generous gift. The human pancreatic islets used in this study were obtained only through the generosity of donor families and their commitment to advancing biomedical research, and this work would not have been possible without their contribution. We honor their legacy by advancing research aimed at improving the understanding and treatment of diabetes and related metabolic diseases. The authors also thank the organ procurement organizations and islet isolation and distribution centers that facilitated the recovery, processing, and distribution of donor pancreatic islets for research purposes.

## Supplementary Material

Supplemental material includes Table S1, the checklist for reporting human islet preparations used in research.

## Data Availability Statement

The data supporting the conclusions of this manuscript will be made available by the authors upon reasonable requests to any qualified researcher.

## Supplementary Material

### Supplementary Tables

**Supplementary Table S1.**
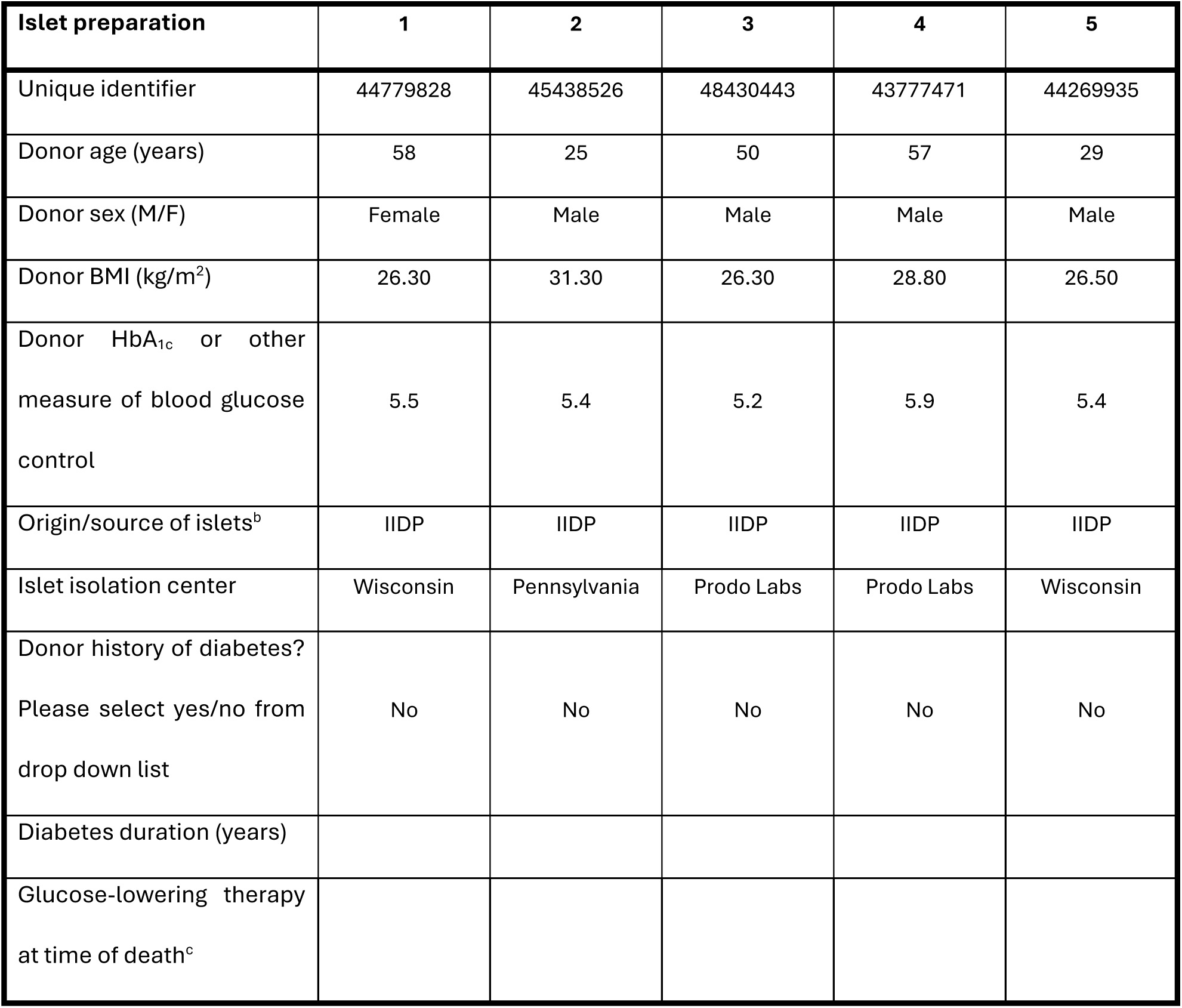
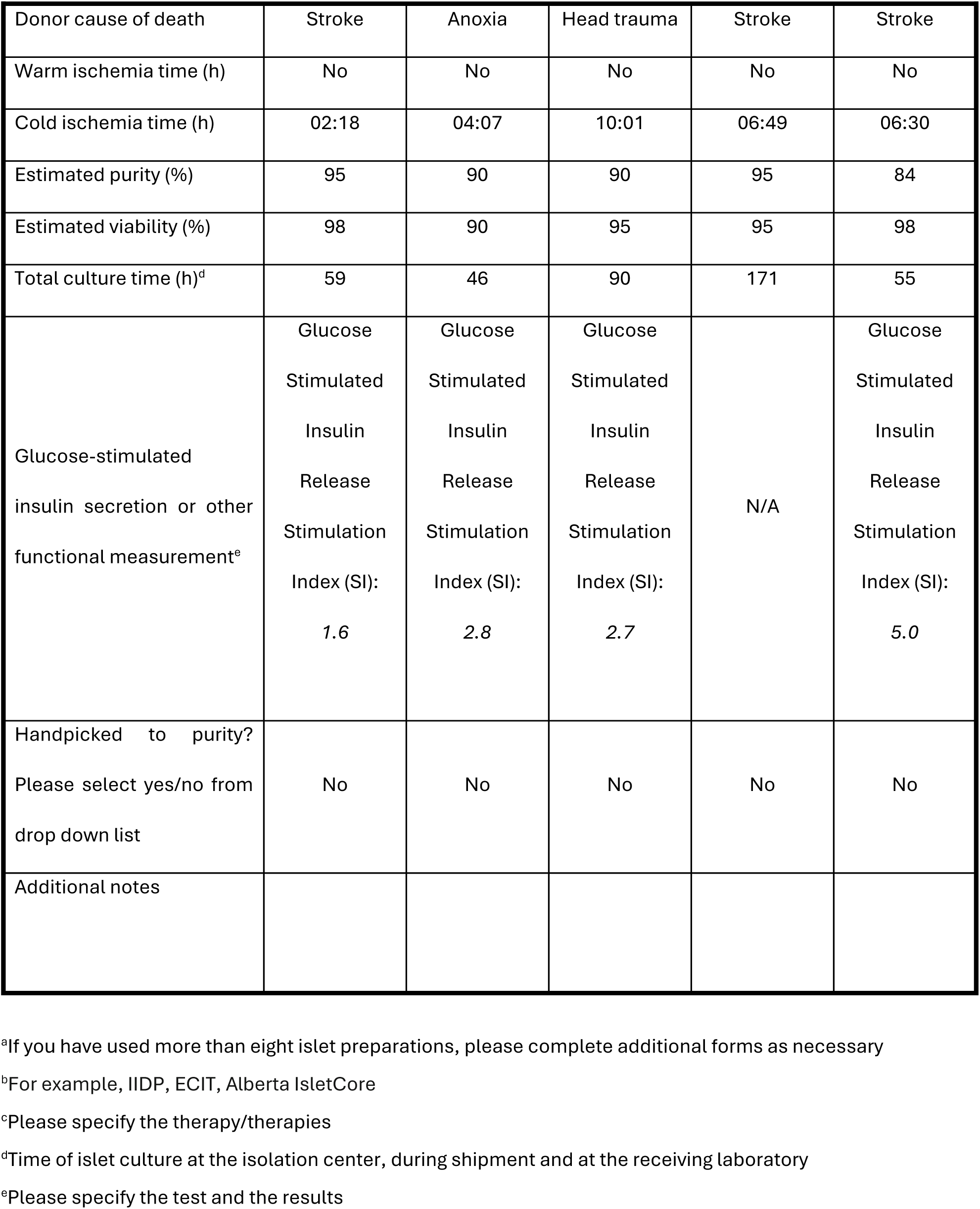
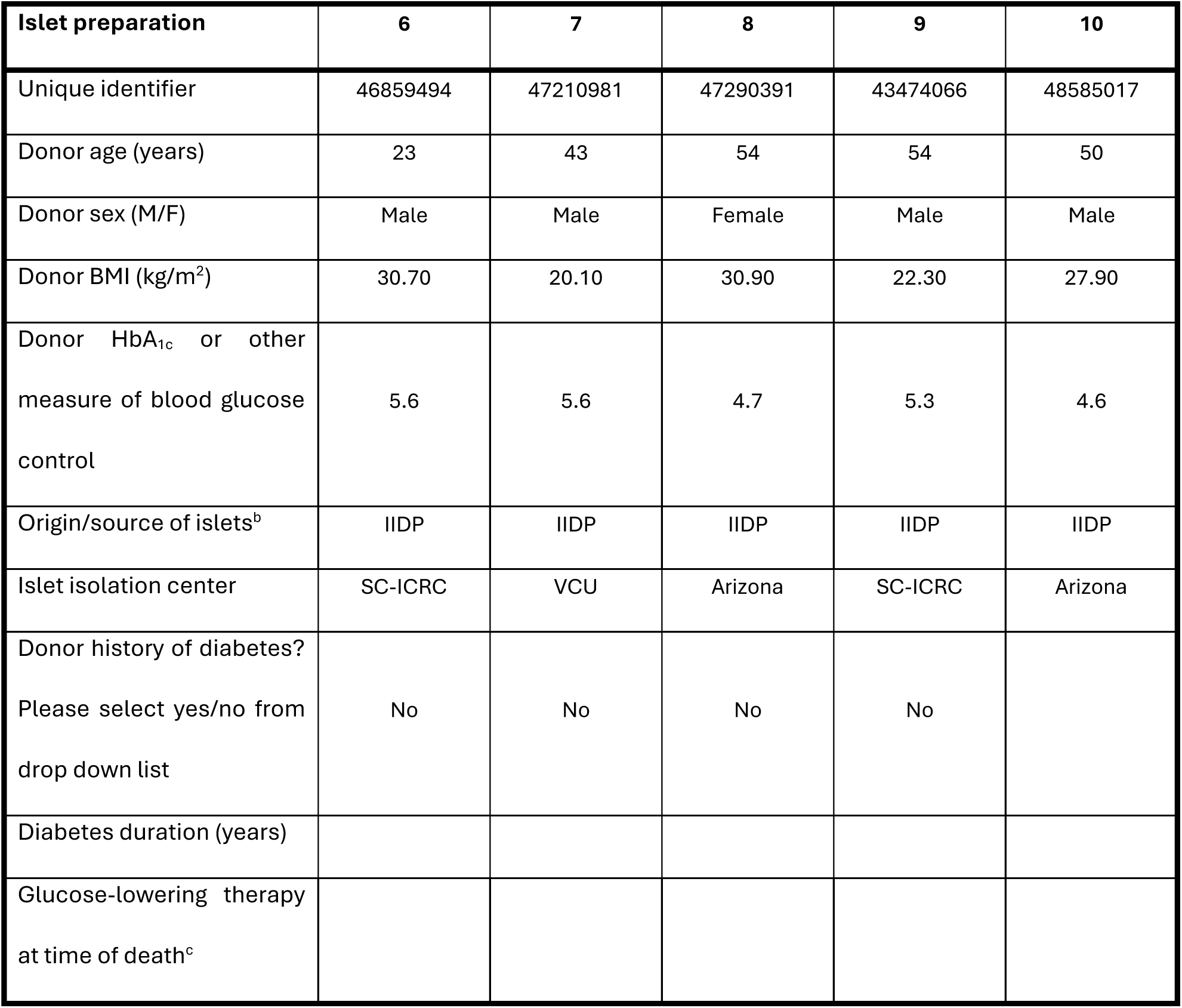
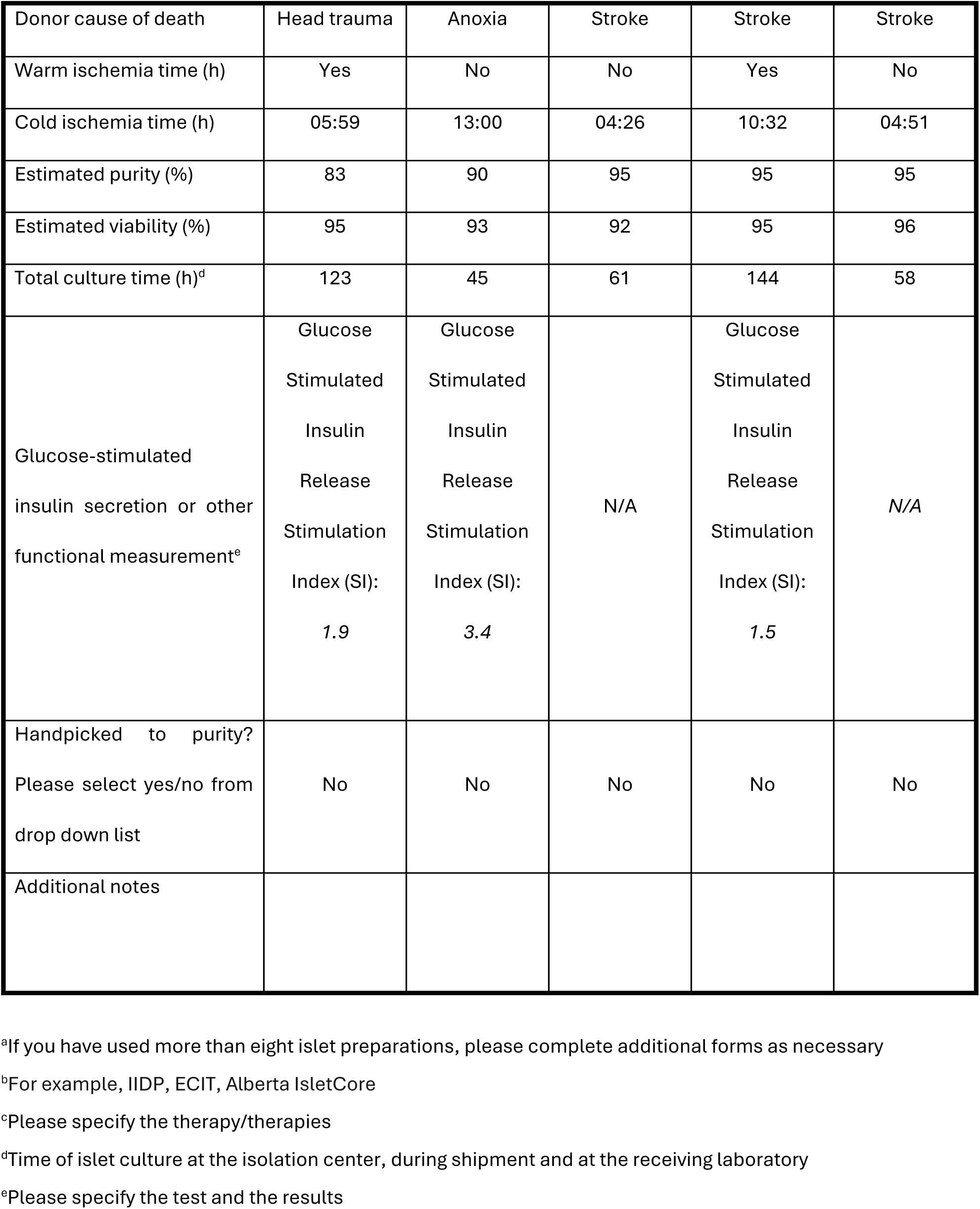
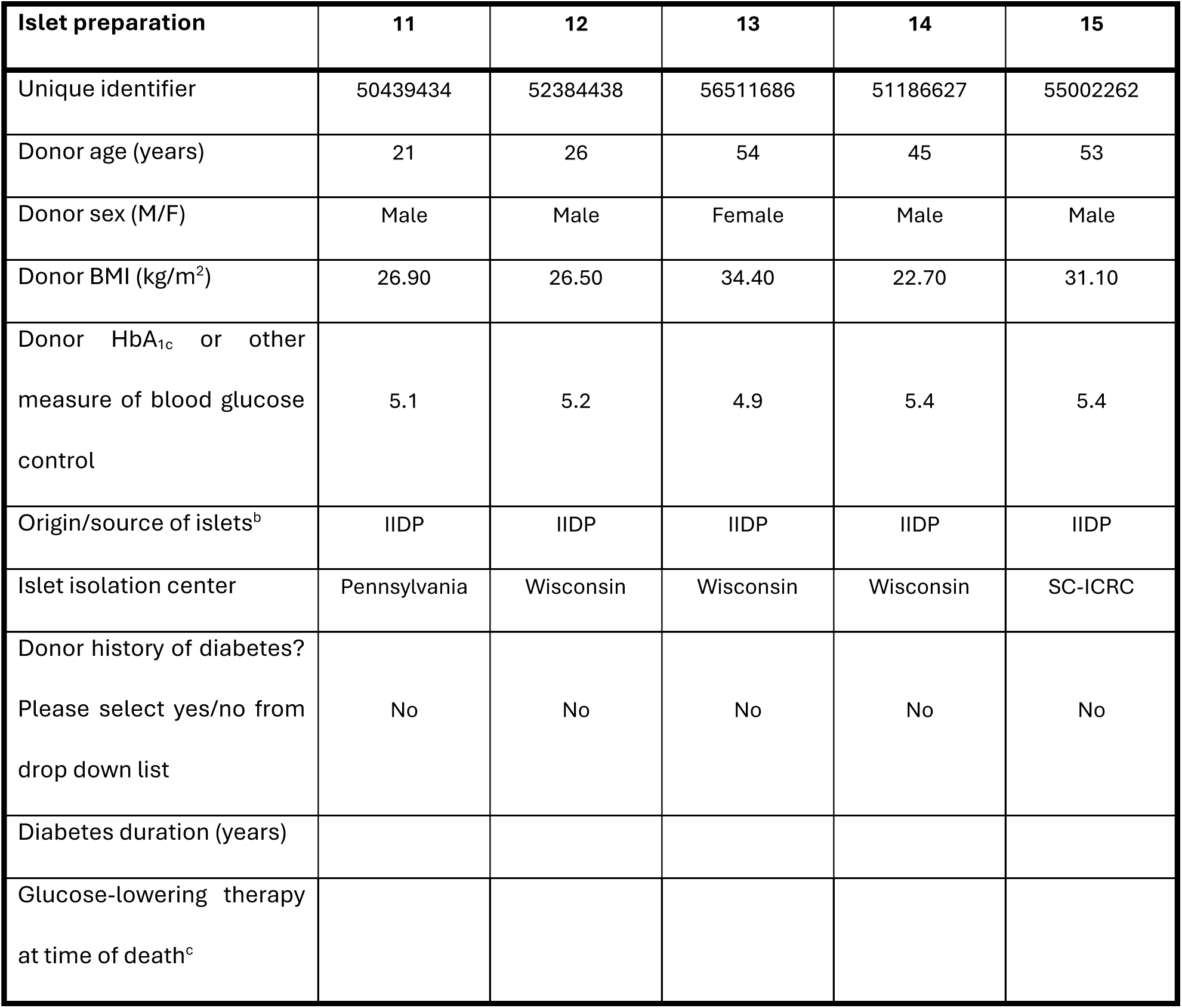
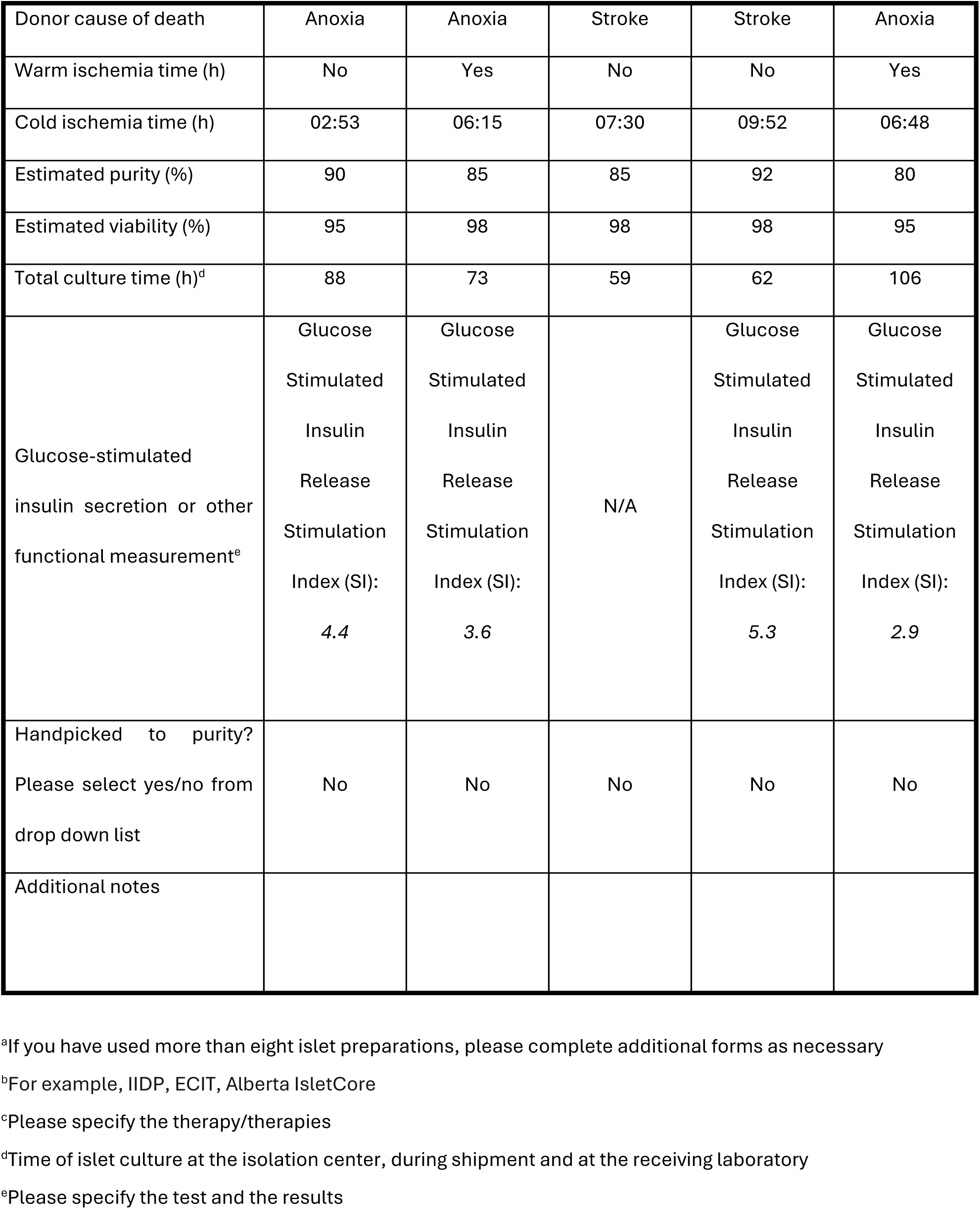
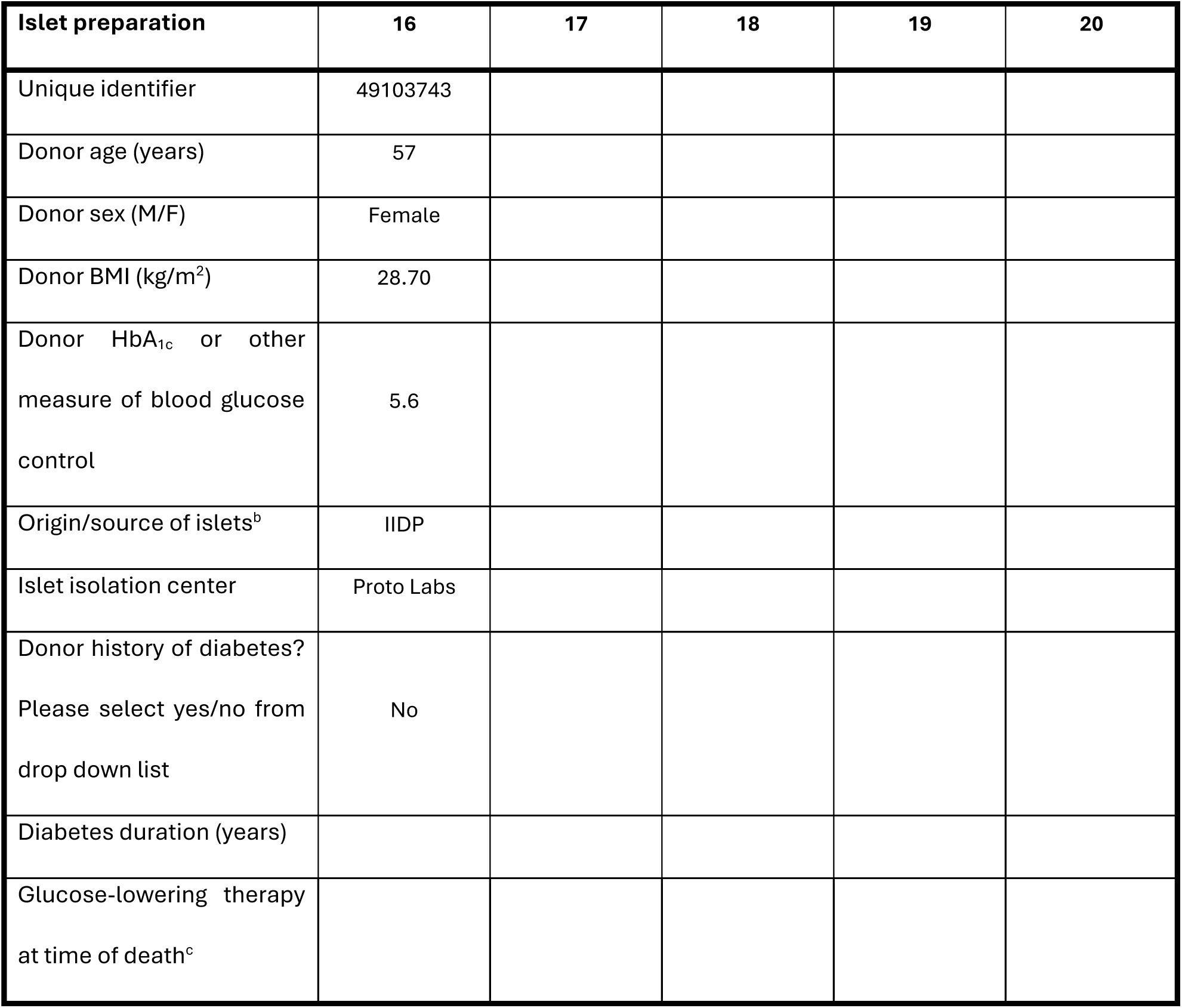
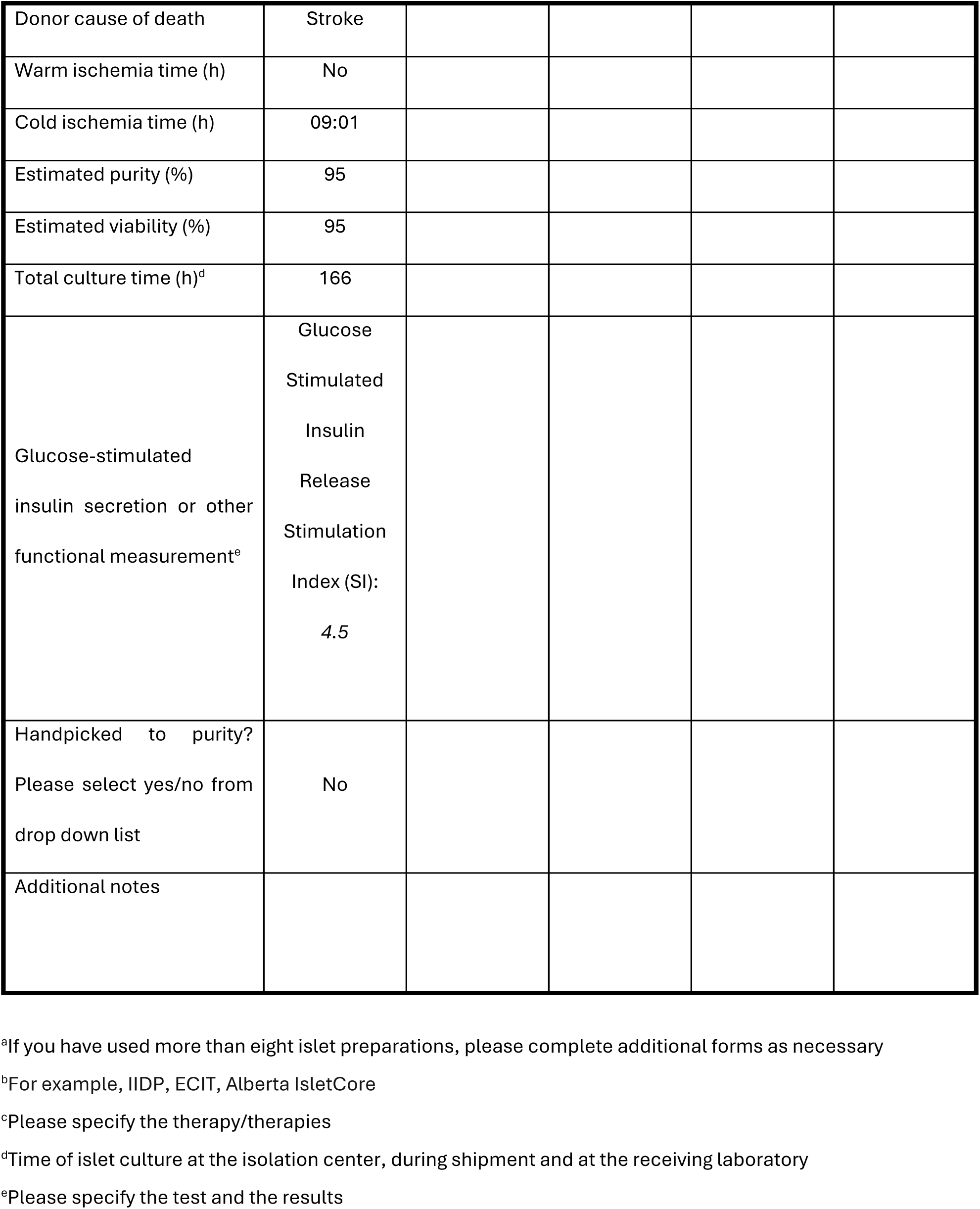
Checklist for Reporting Human Islet Preparations Used in Research. Adapted from Hart NJ, Powers AC (2018) Progress, challenges, and suggestions for using human islets to understand islet biology and human diabetes. *Diabetologia* https://doi.org/10.1007/s00125-018-4772-2.

| Islet preparation | 1 | 2 | 3 | 4 | 5 |
| --- | --- | --- | --- | --- | --- |
| Unique identifier | 44779828 | 45438526 | 48430443 | 43777471 | 44269935 |
| Donor age (years) | 58 | 25 | 50 | 57 | 29 |
| Donor sex (M/F) | Female | Male | Male | Male | Male |
| Donor BMI (kg/m<sup>2</sup>) | 26.30 | 31.30 | 26.30 | 28.80 | 26.50 |
| Donor HbA<sub>1c</sub> or other measure of blood glucose control | 5.5 | 5.4 | 5.2 | 5.9 | 5.4 |
| Origin/source of islets<sup>b</sup> | IIDP | IIDP | IIDP | IIDP | IIDP |
| Islet isolation center | Wisconsin | Pennsylvania | Prodo Labs | Prodo Labs | Wisconsin |
| Donor history of diabetes?<br>Please select yes/no from drop down list | No | No | No | No | No |
| Diabetes duration (years) |  |  |  |  |  |
| Glucose-lowering therapy at time of death<sup>c</sup> |  |  |  |  |  |
| Donor cause of death | Stroke | Anoxia | Head trauma | Stroke | Stroke |
| Warm ischemia time (h) | No | No | No | No | No |
| Cold ischemia time (h) | 02:18 | 04:07 | 10:01 | 06:49 | 06:30 |
| Estimated purity (%) | 95 | 90 | 90 | 95 | 84 |
| Estimated viability (%) | 98 | 90 | 95 | 95 | 98 |
| Total culture time (h)<sup>d</sup> | 59 | 46 | 90 | 171 | 55 |
| Glucose-stimulated insulin secretion or other functional measurement<sup>e</sup> | Glucose<br>Stimulated<br>Insulin<br>Release<br>Stimulation<br>Index (SI):<br>1.6 | Glucose<br>Stimulated<br>Insulin<br>Release<br>Stimulation<br>Index (SI):<br>2.8 | Glucose<br>Stimulated<br>Insulin<br>Release<br>Stimulation<br>Index (SI):<br>2.7 | N/A | Glucose<br>Stimulated<br>Insulin<br>Release<br>Stimulation<br>Index (SI):<br>5.0 |
| Handpicked to purity?<br>Please select yes/no from drop down list | No | No | No | No | No |
| Additional notes |  |  |  |  |  |

<sup>a</sup>If you have used more than eight islet preparations, please complete additional forms as necessary
<sup>b</sup>For example, IIDP, ECIT, Alberta IsletCore
<sup>c</sup>Please specify the therapy/therapies
<sup>d</sup>Time of islet culture at the isolation center, during shipment and at the receiving laboratory
<sup>e</sup>Please specify the test and the results

| Islet preparation | 6 | 7 | 8 | 9 | 10 |
| --- | --- | --- | --- | --- | --- |
| Unique identifier | 46859494 | 47210981 | 47290391 | 43474066 | 48585017 |
| Donor age (years) | 23 | 43 | 54 | 54 | 50 |
| Donor sex (M/F) | Male | Male | Female | Male | Male |
| Donor BMI (kg/m<sup>2</sup>) | 30.70 | 20.10 | 30.90 | 22.30 | 27.90 |
| Donor HbA<sub>1c</sub> or other measure of blood glucose control | 5.6 | 5.6 | 4.7 | 5.3 | 4.6 |
| Origin/source of islets<sup>b</sup> | IIDP | IIDP | IIDP | IIDP | IIDP |
| Islet isolation center | SC-ICRC | VCU | Arizona | SC-ICRC | Arizona |
| Donor history of diabetes?<br>Please select yes/no from drop down list | No | No | No | No |  |
| Diabetes duration (years) |  |  |  |  |  |
| Glucose-lowering therapy at time of death<sup>c</sup> |  |  |  |  |  |
| Donor cause of death | Head trauma | Anoxia | Stroke | Stroke | Stroke |
| Warm ischemia time (h) | Yes | No | No | Yes | No |
| Cold ischemia time (h) | 05:59 | 13:00 | 04:26 | 10:32 | 04:51 |
| Estimated purity (%) | 83 | 90 | 95 | 95 | 95 |
| Estimated viability (%) | 95 | 93 | 92 | 95 | 96 |
| Total culture time (h)<sup>d</sup> | 123 | 45 | 61 | 144 | 58 |
| Glucose-stimulated insulin secretion or other functional measurement<sup>e</sup> | Glucose<br>Stimulated<br>Insulin<br>Release<br>Stimulation<br>Index (SI):<br>1.9 | Glucose<br>Stimulated<br>Insulin<br>Release<br>Stimulation<br>Index (SI):<br>3.4 | N/A | Glucose<br>Stimulated<br>Insulin<br>Release<br>Stimulation<br>Index (SI):<br>1.5 | N/A |
| Handpicked to purity?<br>Please select yes/no from drop down list | No | No | No | No | No |
| Additional notes |  |  |  |  |  |

<sup>a</sup>If you have used more than eight islet preparations, please complete additional forms as necessary
<sup>b</sup>For example, IIDP, ECIT, Alberta IsletCore
<sup>c</sup>Please specify the therapy/therapies
<sup>d</sup>Time of islet culture at the isolation center, during shipment and at the receiving laboratory
<sup>e</sup>Please specify the test and the results

| Islet preparation | 11 | 12 | 13 | 14 | 15 |
| --- | --- | --- | --- | --- | --- |
| Unique identifier | 50439434 | 52384438 | 56511686 | 51186627 | 55002262 |
| Donor age (years) | 21 | 26 | 54 | 45 | 53 |
| Donor sex (M/F) | Male | Male | Female | Male | Male |
| Donor BMI (kg/m<sup>2</sup>) | 26.90 | 26.50 | 34.40 | 22.70 | 31.10 |
| Donor HbA<sub>1c</sub> or other measure of blood glucose control | 5.1 | 5.2 | 4.9 | 5.4 | 5.4 |
| Origin/source of islets<sup>b</sup> | IIDP | IIDP | IIDP | IIDP | IIDP |
| Islet isolation center | Pennsylvania | Wisconsin | Wisconsin | Wisconsin | SC-ICRC |
| Donor history of diabetes?<br>Please select yes/no from drop down list | No | No | No | No | No |
| Diabetes duration (years) |  |  |  |  |  |
| Glucose-lowering therapy at time of death<sup>c</sup> |  |  |  |  |  |
| Donor cause of death | Anoxia | Anoxia | Stroke | Stroke | Anoxia |
| Warm ischemia time (h) | No | Yes | No | No | Yes |
| Cold ischemia time (h) | 02:53 | 06:15 | 07:30 | 09:52 | 06:48 |
| Estimated purity (%) | 90 | 85 | 85 | 92 | 80 |
| Estimated viability (%) | 95 | 98 | 98 | 98 | 95 |
| Total culture time (h)<sup>d</sup> | 88 | 73 | 59 | 62 | 106 |
| Glucose-stimulated insulin secretion or other functional measurement<sup>e</sup> | Glucose<br>Stimulated<br>Insulin<br>Release<br>Stimulation<br>Index (SI):<br>4.4 | Glucose<br>Stimulated<br>Insulin<br>Release<br>Stimulation<br>Index (SI):<br>3.6 | N/A | Glucose<br>Stimulated<br>Insulin<br>Release<br>Stimulation<br>Index (SI):<br>5.3 | Glucose<br>Stimulated<br>Insulin<br>Release<br>Stimulation<br>Index (SI):<br>2.9 |
| Handpicked to purity?<br>Please select yes/no from drop down list | No | No | No | No | No |
| Additional notes |  |  |  |  |  |

<sup>a</sup>If you have used more than eight islet preparations, please complete additional forms as necessary
<sup>b</sup>For example, IIDP, ECIT, Alberta IsletCore
<sup>c</sup>Please specify the therapy/therapies
<sup>d</sup>Time of islet culture at the isolation center, during shipment and at the receiving laboratory
<sup>e</sup>Please specify the test and the results

| Islet preparation | 16 | 17 | 18 | 19 | 20 |
| --- | --- | --- | --- | --- | --- |
| Unique identifier | 49103743 |  |  |  |  |
| Donor age (years) | 57 |  |  |  |  |
| Donor sex (M/F) | Female |  |  |  |  |
| Donor BMI (kg/m<sup>2</sup>) | 28.70 |  |  |  |  |
| Donor HbA<sub>1c</sub> or other<br>measure of blood glucose<br>control | 5.6 |  |  |  |  |
| Origin/source of islets<sup>b</sup> | IIDP |  |  |  |  |
| Islet isolation center | Proto Labs |  |  |  |  |
| Donor history of diabetes?<br>Please select yes/no from<br>drop down list | No |  |  |  |  |
| Diabetes duration (years) |  |  |  |  |  |
| Glucose-lowering therapy<br>at time of death<sup>c</sup> |  |  |  |  |  |

|  |  |
| --- | --- |
| Donor cause of death | Stroke |
| Warm ischemia time (h) | No |
| Cold ischemia time (h) | 09:01 |
| Estimated purity (%) | 95 |
| Estimated viability (%) | 95 |
| Total culture time (h)<sup>d</sup> | 166 |
| Glucose-stimulated insulin secretion or other functional measurement<sup>e</sup> | Glucose<br>Stimulated<br>Insulin<br>Release<br>Stimulation<br>Index (SI):<br>4.5 |
| Handpicked to purity?<br>Please select yes/no from drop down list | No |
| Additional notes |  |

<sup>a</sup>If you have used more than eight islet preparations, please complete additional forms as necessary
<sup>b</sup>For example, IIDP, ECIT, Alberta IsletCore
<sup>c</sup>Please specify the therapy/therapies
<sup>d</sup>Time of islet culture at the isolation center, during shipment and at the receiving laboratory
<sup>e</sup>Please specify the test and the results

